# PodoCount: A robust, fully automated whole-slide podocyte quantification tool

**DOI:** 10.1101/2021.04.27.441689

**Authors:** Briana A. Santo, Darshana Govind, Parnaz Daneshpajouhnejad, Xiaoping Yang, Xiaoxin X. Wang, Komuraiah Myakala, Bryce A. Jones, Moshe Levi, Jeffrey B. Kopp, Laura J. Niedernhofer, David Manthey, Kyung Chul Moon, Seung Seok Han, Avi Z. Rosenberg, Pinaki Sarder

**Affiliations:** Department of Pathology and Anatomical Sciences, University at Buffalo, Buffalo, NY; Department of Pathology, Johns Hopkins University School of Medicine, Baltimore, MD; Department of Biochemistry, Molecular & Cellular Biology, Georgetown University, Washington, DC; Department of Pharmacology and Physiology, Georgetown University, Washington, DC; Kidney Disease Section, NIDDK, NIH. Bethesda, MD; Institute on the Biology of Aging and Metabolism, Department of Biochemistry, Molecular Biology, and Biophysics, University of Minnesota, Minneapolis, MN; Kitware, Clifton Park, NY; Department of Pathology, Seoul National University College of Medicine, Seoul, Korea; Department of Internal Medicine, Seoul National University College of Medicine, Seoul, Korea

**Keywords:** Podocyte, podometrics, glomerulus, glomerular disease, chronic kidney disease, digital pathology, gigapixel size images

## Abstract

**Background:** Podocyte depletion is an established indicator of glomerular injury and predicts clinical outcomes. The semi-quantitative nature of existing podocyte estimation methods or podometrics hinders incorporation of such analysis into experimental and clinical pathologic workflows. Computational image analysis offers a robust approach to automate podometrics through objective quantification of cell and tissue structure. Toward this goal, we developed PodoCount, a computational tool for quantitative analysis of podocytes, and validated the generalizability of the tool across a diverse dataset.

**Methods:** Podocyte nuclei and glomerular boundaries were labeled in murine whole kidney sections, *n* = 135, from six disease models and human kidney biopsies, *n* = 45, from diabetic nephropathy (DN) patients. Digital whole slide images (WSIs) of tissues were then acquired. Classical image analysis was applied to obtain podocyte nuclear and glomerular morphometrics. Statistically significant morphometric features, which correlated with each murine disease, were identified. Engineered features were also assessed for their ability to predict outcomes in human DN. *PodoCount* has been disbursed for other researchers as an open-source, cloud-based computational tool.

**Results:** *PodoCount* offers highly accurate quantification of podocytes. Engineered podometric features were benchmarked against routine glomerular histopathology and were found to be significant predictors of disease diagnosis, proteinuria level, and clinical outcomes.

**Conclusions:** *PodoCount* offers high quantification performance in diverse murine disease models as well as in human DN. Resultant podometric features offers significant correlation with associated metadata as well as outcome. Our cloud-based end-user tool will provide a standardized approach for podometric analysis from gigapixel size WSIs in basic research and clinical practice.

## 1. INTRODUCTION

Chronic kidney disease (CKD) is a state of reduced kidney function that may progress to end-stage kidney disease (ESKD). Driven by increasingly prevalent conditions with high incidence (e.g., diabetes, hypertension), CKD will account for >1.2 million deaths and unprecedented socioeconomic burden in 2021^1,2^. To mitigate this, biomedical initiatives aim to identify disease-related biomarkers with improved precision for early detection and intervention. Current studies look to animal models and human biopsies for guidance on prospective biomarkers. One such example is the assessment of highly specialized epithelial cells, podocytes, that form part of the glomerular filtration barrier (GFB). According to the podocyte depletion hypothesis^3–6^, injury-driven loss of podocytes contributes to glomerular injury and thus CKD progression. A myriad of studies confirms podocyte depletion as an early determinant of proteinuria and glomerulosclerosis^7–9^. These observations rendered podocytes a measurable indicator of renal injury and therapeutic success in CKD.

Unfortunately, podocyte analysis in experimental and clinical workflows is limited to semi-quantitative methods of podocyte estimation that poorly approximate whole-slide cell counts, provide little-to-no morphological assessment, and overlook regional variation in podocyte pathology^3,4,10–13^. Further complicating this situation is the fact that podocyte identification on routine and special stains viewed under brightfield histopathology remains difficult ^10^. In order to achieve big-data podocyte studies that facilitate early detection and intervention in basic research and clinical practice, accurate and precise quantitative methods must be developed to identify, characterize and contextualize podocytes in brightfield microscopy.

Computational image analysis offers a unique approach to refine podocyte estimation through objective quantification of cell and tissue structures in digital pathology. In their review*, Hodgin, Wiggins et al*. highlight how automated podometrics will augment podocyte-centric clinical studies and basic research ^5,14,15^. Podometrics is defined as a set of techniques for quantification of well-validated podocyte metrics, including the quantification of glomerular podocyte (nuclear) number, size, and density from immunohistochemistry (IHC)-labeled kidney biopsies^4,5,10–14,16^. Toward this goal, we developed *PodoCount* – a tool for automated podometrics in digitized kidney biopsies (***Fig 1***). Our work is enabled by recent advancements in whole-slide imaging and hardware technologies^17,18^, as well as an optimal staining technique that labels podocyte nuclei and glomerulus boundaries in tissue specimens.

**Figure 1:**
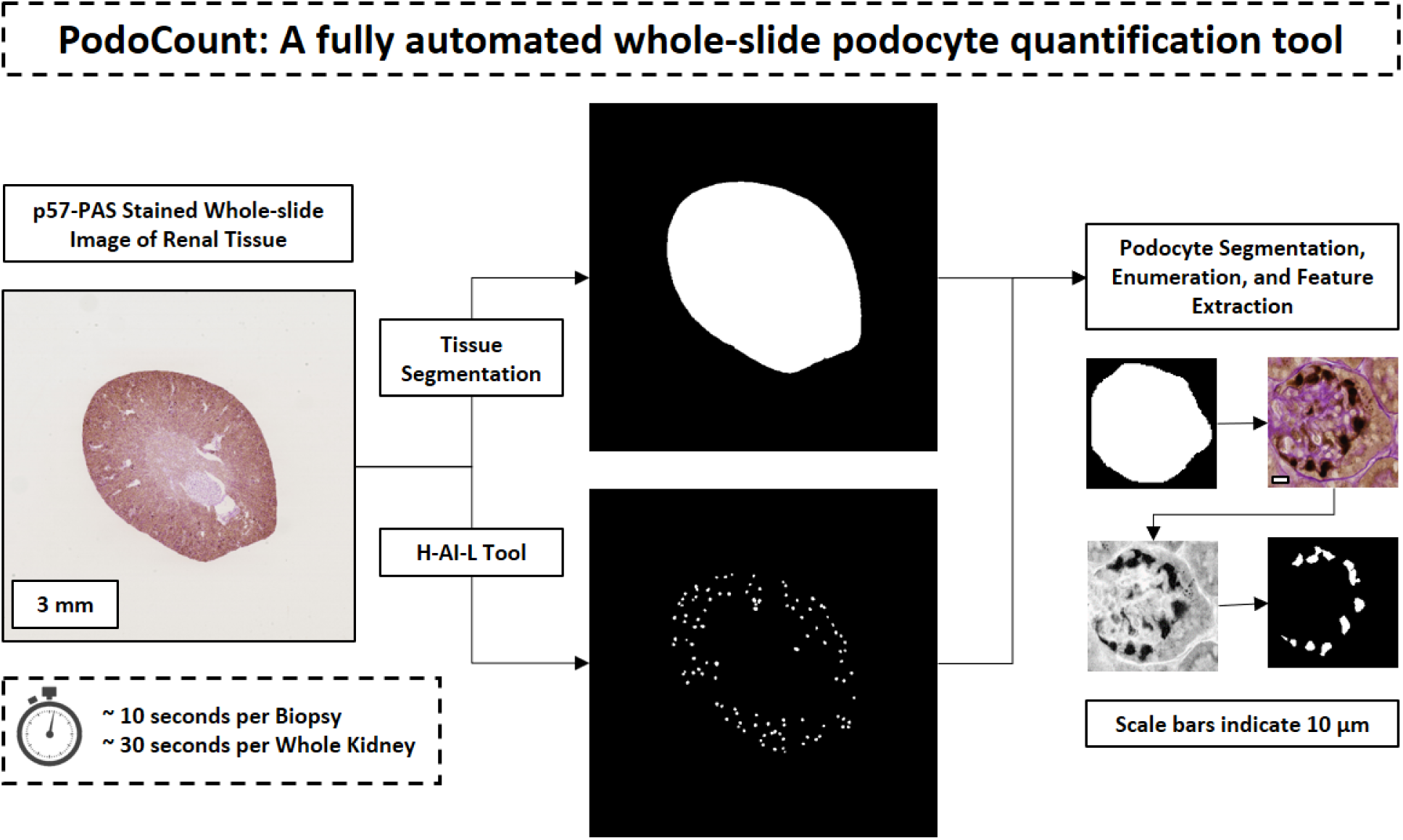
A robust tool for whole-slide podocyte quantification was developed. *Legend.* Shown is a flow-chart of the image analysis pipeline for quantification of podocyte depletion in immunohistochemically-labeled kidney specimen. Whole slide images (WSI) of kidney tissues and corresponding glomerulus annotation files are entered into a dedicated informatics pipeline, which segments podocytes, glomeruli, and tissue sections from WSI in order to (i) enumerate podocytes and (ii) obtain podocyte and glomerulus morphometrics.

To validate our automated podometrics, we applied *PodoCount* in a multi-institutional and multi-species dataset comprising six glomerular disease models and human diabetic nephropathy (DN). This dataset emulates the heterogeneity of disease pathology, sample preparation, and digitization characteristic of digital pathology data^19–23^. Computational evaluation demonstrated that *PodoCount* achieved high performance in quantifying podocyte nuclei. Subsequent statistical analysis suggested that computed podocyte metrics highlight subtle differences in histopathology essential for differentiation of wild-type and disease states as well as critical stages in DN progression. To ensure our pipeline’s broad applicability and accessibility, we optimized *PodoCount* across this vast dataset and developed a web-based plugin for podometrics in the cloud. Cloud-based podocyte counting provides a user-friendly, universally accessible environment for computational analysis irrespective of one’s programming experience or operating system. Operating at an efficient rate and high performance, *PodoCount* has tremendous potential to ease podocyte-centric experimental and clinical workflows.

## 2. METHODS

### 2.1 Disclosure

Human data collection followed protocols approved by the Institutional Review Board at the Seoul National University (SNU) College of Medicine (H-1812-159-998), Seoul, Korea. All experiments were performed according to federal guidelines and regulations. Animal studies were performed in accordance with protocols approved by the Institutional Animal Care and Use Committee at the Georgetown University, National Institutes of Health, University of Minnesota, and Johns Hopkins University (JHU), are consistent with federal guidelines and regulations, and are in accordance with recommendations of the American Veterinary Medical Association guidelines on euthanasia.

### 2.2 Murine and Human data

This study used data from six murine kidney disease models and kidney biopsies from a diabetic nephropathy patient cohort. A brief summary of each dataset has been provided below (***Fig 2***). These multi-institutional, multi-species specimens, of both male and female origin, feature highly variable sample preparation, imaging, staining and pathology, and thus comprise a dataset that exemplifies scientific rigor and reproducibility (***Fig 3***).

**Figure 2:**
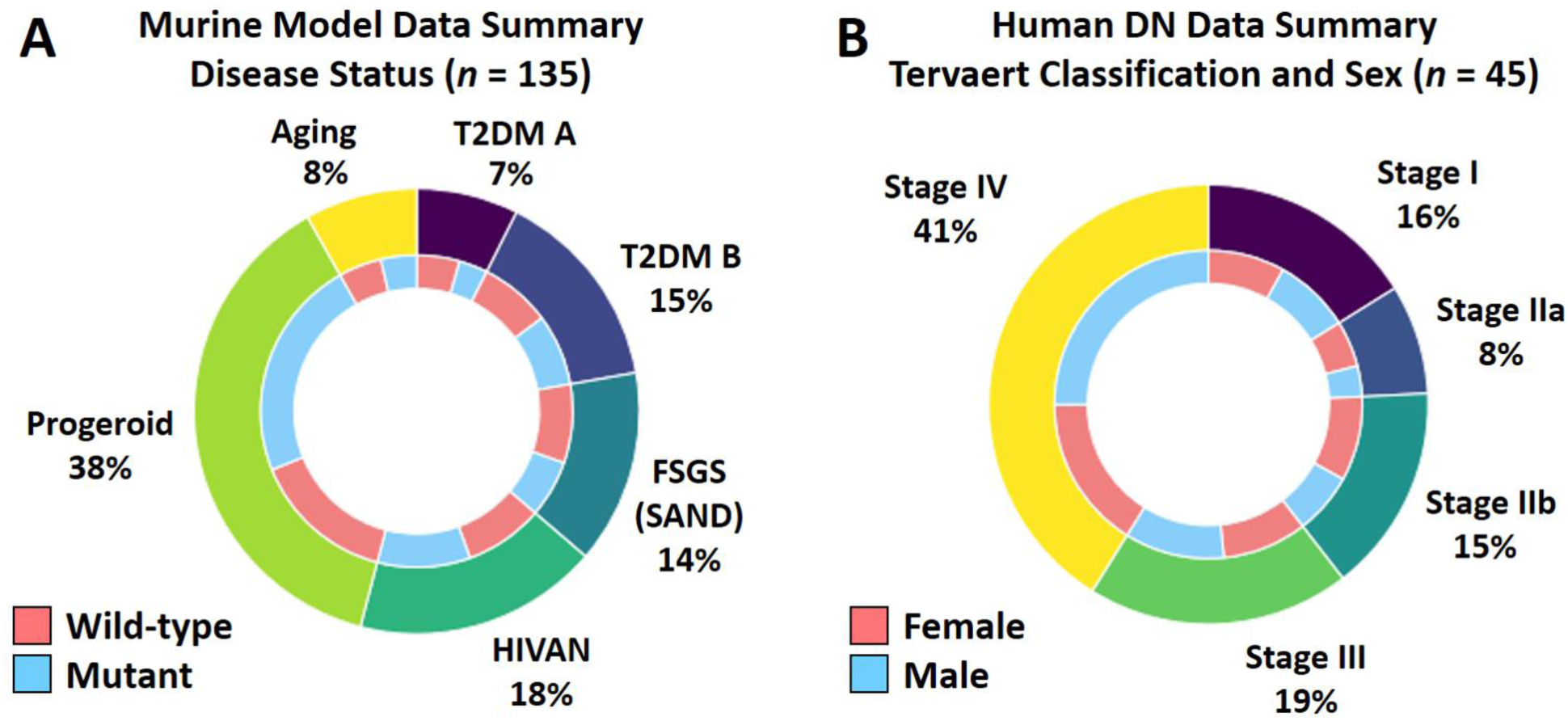
A rigorous dataset was used to develop and validate our method. T2DM, type-II diabetes mellitus; FSGS (SAND), a post-adaptive model of FSGS, focal segmental glomerulosclerosis; HIVAN, human immunodeficiency virus-associated nephropathy. *Legend.* The image dataset contains light-microscopic images of kidney tissues from six mouse models of glomerular disease and five stages of human diabetic nephropathy (DN). Shown are the numbers of wild-type and mutant or disease-induced mice. The murine cohort totaled 135 samples and included both normal and diseased mice. Two distinct models of type-II diabetes mellitus were studied and are denoted as T2DM A and T2DM B. The SAND intervention (saline, angiotensin II, uninephrectomy, and deoxycortisone) models post-adaptive FSGS, and thus the model is termed FSGS (SAND). The mouse SAND, HIVAN, and Progeroid syndrome studies include both males and females, while the T2DM A, T2DM B, and Aging mouse models included only males. The human DN study included both male and female subjects.

**Figure 3:**
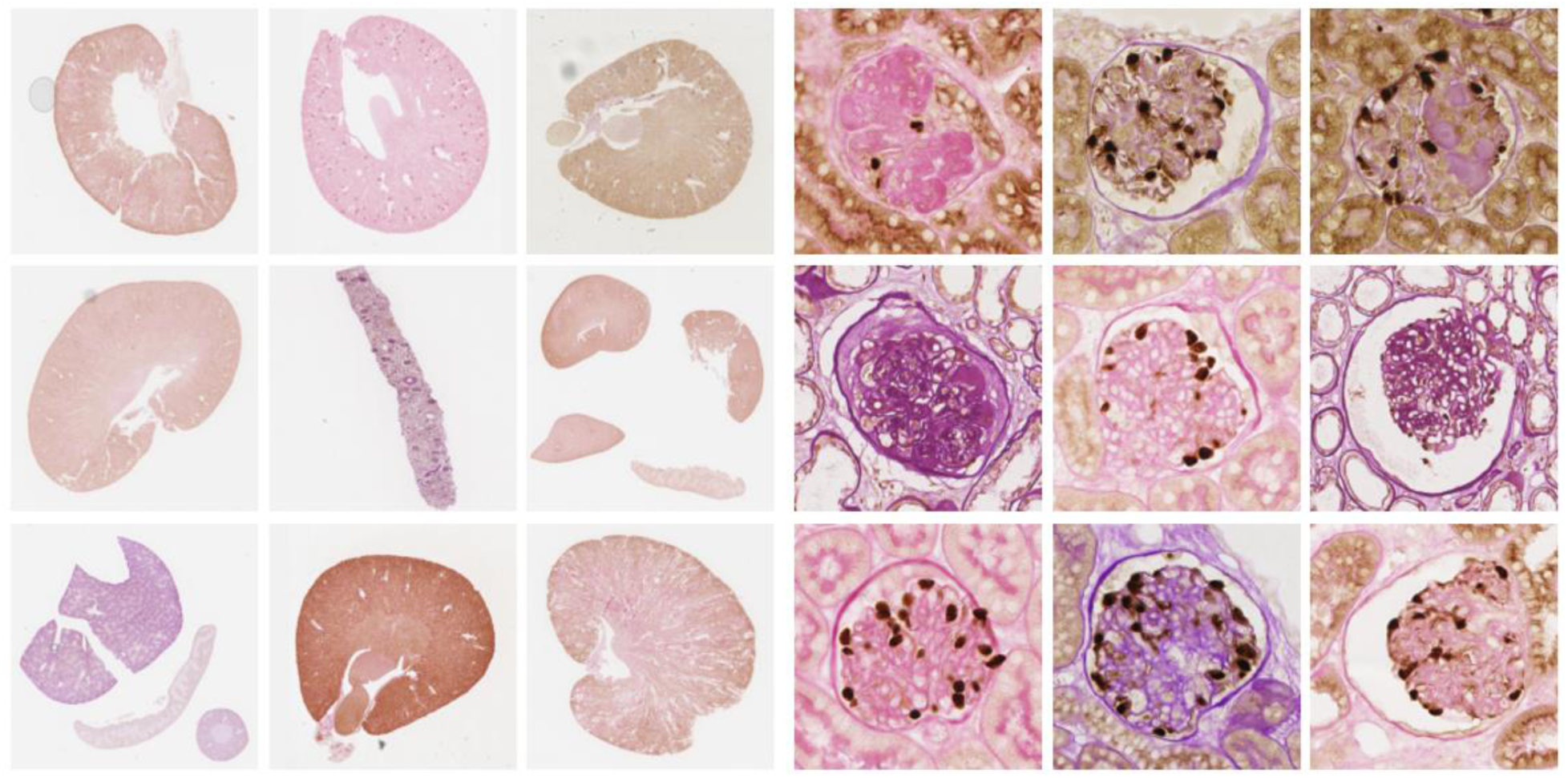
*PodoCount* manifests scientific rigor and reproducibility. *Legend.* PodoCount is a generalizable computational tool for podocyte quantification. This tool successfully assessed multi-institutional and multi-species data featuring highly variable sample preparation, imaging, and staining, and diverse histopathology. This dataset exemplifies the inherently challenging nature of digital pathology data, which drives the need for generalizable computational frameworks.

*Murine cohort: Cohort 1:* An HIV-associated nephropathy (HIVAN) model was used. In this model, transgenic Tg26 mice on a FVB/N background feature a *gag-pol*-deleted HIV-1 genome^24^ which manifests as collapsing glomerulopathy. *Cohort 2:* Wild-type FVB/N mice were subjected to a combination of four interventions that induce a post-adaptive form of FSGS, summarized by the acronym SAND. The interventional process includes 0.9% saline drinking water, angiotensin II infused via osmotic pump, uni-nephrectomy, and deoxycorticosterone delivered by implantation of a subcutaneous pellet.^25,26^ Cohort 2 is referred to as FSGS (SAND) throughout this study. *Cohort 3 & 4:* Two models of type-2 diabetes mellitus (T2DM) were used in this study, summarized as T2DM A and T2DM B throughout this work. For T2DM A (*Cohort 3*), db/db mice on BKS background featuring a leptin receptor mutation were used. These mice depict spontaneous/congenital diabetes due to leptin signaling abnormalities^27^. For T2DM B (*Cohort 4*), the KKAy mouse model (see description in a previous work^28^) was used that develop spontaneous diabetes of polygenic origin. *Cohort 5:* Aging studies were performed in 4-month-old and 21- month-old C57BL/6 male mice obtained from the NIA aging rodent colony^29^. *Cohort 6:* An *Ercc1^-/Δ^* Progeroid mouse model and wild-type littermate controls (15-18-week-old) on a C57BL/6J:FVB/N f1 background were used for the study^30,31^. Mice were bred and genotyped as previously described^32^.

*Human cohort:* Human tissues consisted of needle biopsy samples from human T2DM patients (*n* = 45). Samples were collected from the Seoul National University Hospital Human Biobank. Biopsies were graded by a renal pathologist based on the Tervaert classification method^33^. Clinical metadata including estimate glomerular filtration rate (eGFR)^34^ and serum creatinine were measured during biopsy and one-year and two-years post biopsy. This study considered progression to ESKD within 2 years following biopsy as the main outcome.

### 2.3 Sample preparation and imaging

Sample preparation and staining procedures were consistent with existing protocols for IHC staining of formalin fixed, paraffin embedded (FFPE) tissues. Podocyte nuclei were immunohistochemically labeled in tissue sections using an antibody specific for p57^kip2^, a marker of podocyte terminal differentiation^35^ (primary ab75974, Abcam, Cambridge, UK), followed by HRP Unovue Rabbit HRP detection reagent (RU-HRP1000, Diagnostic BioSystems, Pleasanton, CA) and exposed to diaminobenzidine (DAB) chromogen and substrate (BSB0018A, Bio SB, Santa Barbara, CA). A periodic acid-Schiff (PAS) post-stain was applied. Hematoxylin counterstain was omitted from the traditional PAS protocol.

Bright-field whole slide images (WSI) were captured using an Aperio AT2 microscope (Leica Microsystems, Buffalo Grove, Il) or a NanoZoomer S360 slide scanner equipped with a 40X objective (Hamamatsu Photonics, Bridgewater, NJ). The full image dataset consisted of WSI of *n* = 135 whole murine kidney sections and *n* = 45 DN biopsies (Fig 2).

### 2.4 Whole slide compartmentalization of renal parenchyma

Images of podocytes and renal tissue compartments were extracted from WSI through image segmentation by selecting image regions of interest (ROIs) based on differences in color, texture, and shape^36^. Segmented structures included whole tissue sections, glomerular boundaries, and podocyte nuclei.

Structural segmentation involved several steps. For whole tissue sections, first a representative color vector derived from the original red-green-blue (RGB) WSI was applied as a global mean-based threshold to segment the tissue section from the background of the WSI. Then glomerular boundaries were detected using our previously published Human-AI-Loop (H-AI-L) tool^37^, which is a convolutional neural network developed for WSI segmentation of digital pathology. In the next step, podocyte nuclei were segmented from each detected glomerulus. Toward this goal, first a stain deconvolution algorithm^38^ was used to separate the IHC-positive (IHC+) podocyte nuclei from the surrounding PAS-positive (PAS+) glomerulus micro-compartments. A local-mean based threshold was applied to produce a binary image, with podocyte nuclei in the foreground. Morphological image processing techniques^36^ including hole-filling and size exclusion were applied to remove excess nuclear DAB, while marker controlled watershed^39–41^ separated overlapping nuclei. In image processing, a watershed transformation involves analyzing the image as if it were a topographic map, with image intensity being represented as height on a map, and the addition of lines that separate regions of similar image intensity. A marker representing the peak of each region may be superimposed on the map to control or guide the placement of these separation lines for optimal division of contiguous regions.

### 2.5 Quantitative performance analysis

Sensitivity, specificity, precision, and accuracy were computed in order to evaluate pipeline segmentation performance. Podocyte counts were evaluated by comparison with manual counts. Performance analysis was conducted for a subset of WSIs (*n* = 12, including both wild-type and mutant mice from each model) and corresponding glomerulus ROIs (*n* = 120) in order to assess the accuracy of segmentation and enumeration of podocytes. For all images, podocyte nuclei and tissue and glomerulus boundaries, were annotated manually to provide ground truth for comparison with automatic segmentations. In addition, podocytes were manually counted in each extracted glomerulus image for comparison with pipeline counts. For each of the twelve WSIs, segmentation and enumeration metrics were computed across all ROIs, and subsequently were computed for all WSIs.

### 2.6 Whole-slide podocyte and glomerulus feature extraction

Segmented podocyte nuclei were enumerated in each ROI as an estimation of glomerular podocyte count. Built-in morphological operations were applied to derive geometric features from both podocytes and glomeruli^42^. Geometric features included image object area, bounding box area, convex area, eccentricity, equivalent diameter, extent, major and minor axis lengths, orientations, perimeters, and solidities. A brief description of each feature is provided in ***Table S1***. Pixel intensity statistics were derived from podocyte nuclei exclusively. In addition to individual feature values, feature statistics were computed for each single-glomerulus podocyte population as well as for the whole-slide glomerulus population. All morphometric features were recorded in csv file format for subsequent statistical analysis.

### 2.7 Biologically-inspired podocyte feature engineering

To best measure podocyte depletion (***Table 1***), features were engineered to emulate podocyte pathology in glomerular disease. For each glomerulus unit, podocyte spatial density was estimated using three steps: (i) dividing the absolute podocyte count by the glomerular area, (ii) computing the total area of podocyte nuclei within the glomerulus ROI, and (iii) dividing the cumulative podocyte nuclear area by the glomerular area. These area-based metrics were converted to µm^2^ in order to place measurements within familiar spatial context. Feature values were recorded for subsequent analysis.

**Table 1.**
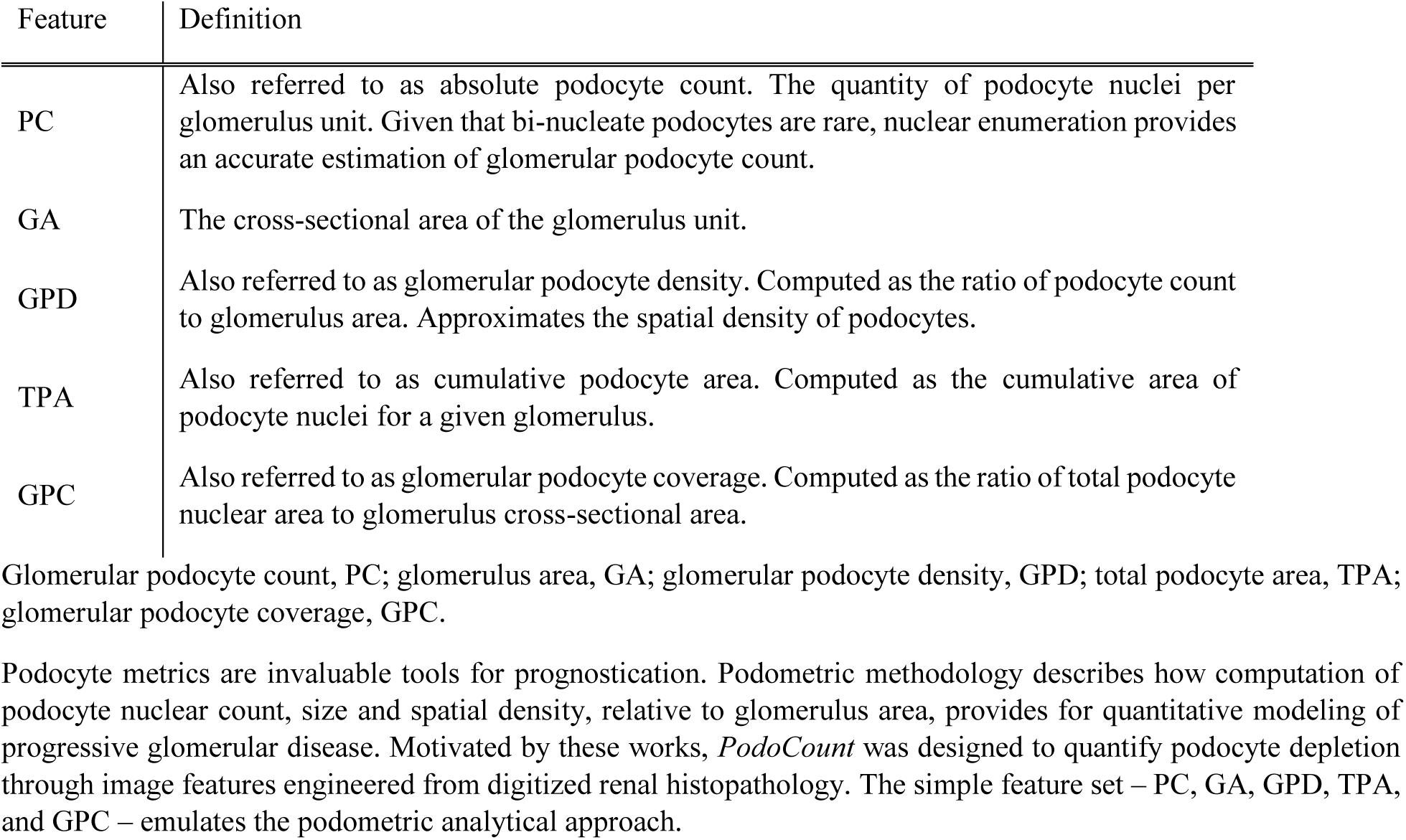
Histologic image features were selected for statistical analysis based on established Podometrics.

### 2.8 Statistical analysis

Data were analyzed with Minitab Statistical Software v19 [Minitab 17 Statistical Software (2010), Minitab, State College, PA]. All analysis was completed at a significance level (*α*) of 0.05.

Differences between groups, including, wild-type and mutant mice and male vs female mice, were assessed with unpaired, two-sample *t*-tests. Welch’s correction for unequal variances was performed when appropriate. Correlation analysis between murine histologic image features and urinary albumin/creatinine ratio (UACR) at time of sacrifice were completed using the Pearson correlation coefficient measure (Pearson’s *R*)^43^.

Differences among three or more groups, including diabetic nephropathy stages, were assessed with one-way ANOVA, followed by *post hoc* Bonferroni tests for multiple comparisons. For those feature distributions which violated ANOVA criteria, the Kruskal-Wallis nonparametric test was used to compare population medians, followed by *post hoc* Dunn’s tests^44,45^. Logistic regression was used to study binary ESKD outcome, with a chi-square test comparing the simple linear regression against a null model^46^.

### 2.9 Deployment of whole-slide podocyte analysis with cloud computation

HistomicsUI^47^, a distributed system with RESTful application programming interface (API), was developed by Kitware (Clifton Park, NY) and was used to deploy our algorithm as a plugin, thereby creating an online platform which would enable multiple users to detect and quantify podocytes via a web interface. The algorithm was packaged in the form of a Docker image using *Docker* softwar*e* (Palo Alto, CA) ^48–50^, a framework that enables users to build and run applications in containers. The generated container conforms to the Slicer CLI workflow interface, which allows *HistomicsUI* to display a user interface to adjust algorithm parameters.

### 2.10 Hardware

Computational processing was performed on a Linux distribution operating system (Ubuntu 16.04) with two Intel Xeon Silver 4114 processors, each with 10 cores, running at 2.20 GHz and equipped with 64 GB of RAM. Neural network training and predictions for glomerulus boundary detection were performed using a NVIDIA Quadro RTX 5000 GPU (16 GB of memory). *HistomicsUI* plugin is available in a standard research computer without any GPU with Intel i5 6-core processor, running at 3.1 – 4.5 GHz and equipped with 16 GB of RAM.

### 2.11 Data availability

To support reproducibility, we released fully-annotated pipeline codes for other researchers to use, along with WSIs and pipeline outputs. We also launched our cloud-based *PodoCount* plugin for the end-user community. The plugin link, all codes and comprehensive documentation, as well as the docker image of the web cloud interface, select WSIs, segmented outputs, and quantified feature files will all be made publicly available following manuscript publication.

## 3. RESULTS

### 3.1 Qualitative performance analysis

Visual inspection of pipeline-derived podocyte nuclear, glomerular, and tissue boundaries confirmed successful region detection as well as segmentation (***Fig 4***). Automated podometrics requires high-fidelity separation of closely spaced, overlapping podocyte nuclei, and observed segmentation results underscore the utility of a marker-controlled watershed for our nuclear segmentation task.

**Figure 4:**
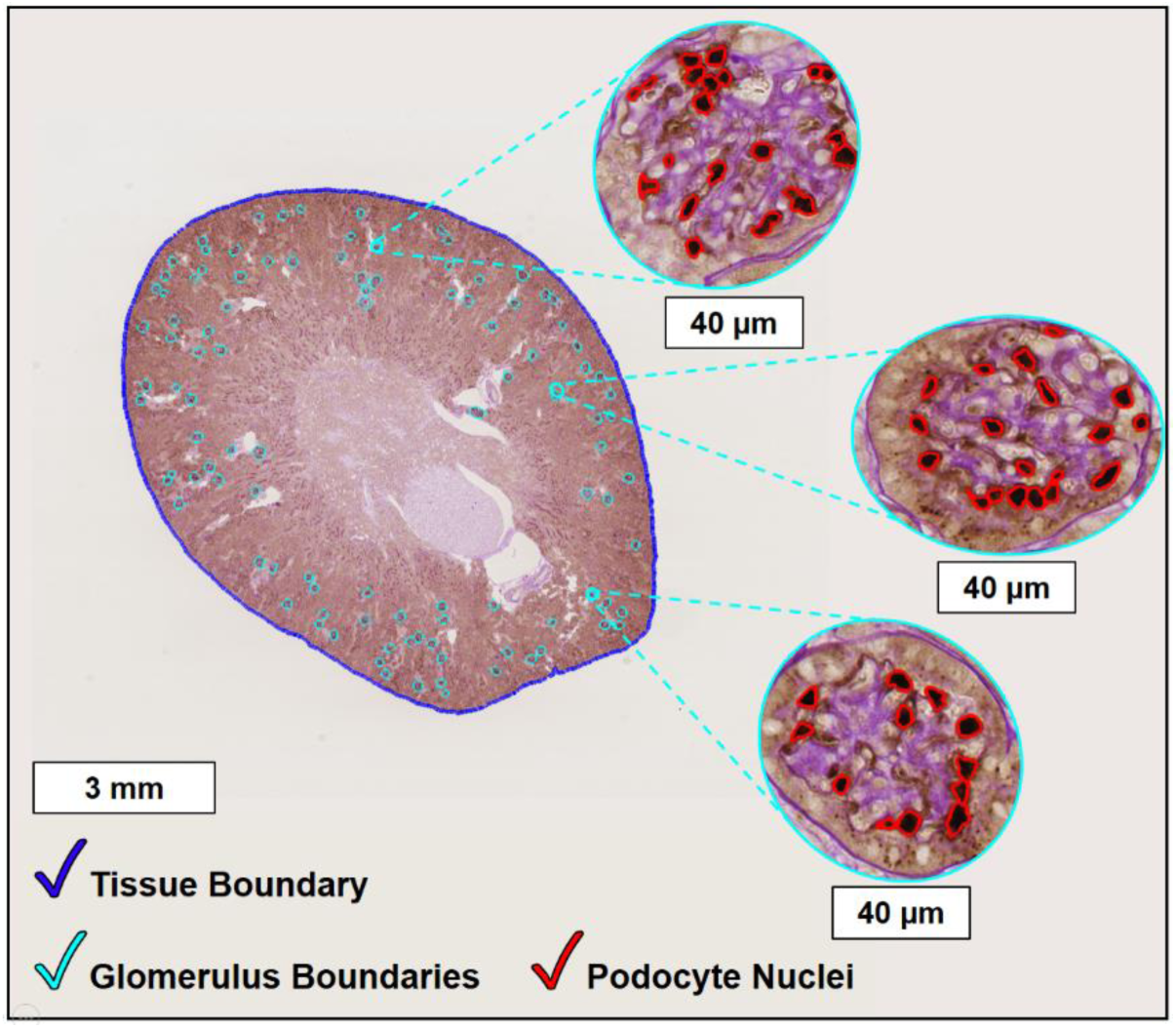
Whole-slide image segmentation enabled quantification of nuclear and tissue structure. *Legend.* Classical image analysis techniques were applied whole-slide images, in order to segment tissue boundaries (dark blue) and identify immunohistochemically-labeled podocyte nuclei (red). Glomerulus boundaries (cyan) were detected using the H-AI-L tool (Lutnick *et al*., *Nature Machine Intelligence,* 2019) which is a convolutional neural network for glomerular boundary detection.

### 3.2 Quantitative performance analysis

The median performance across all twelve WSIs was computed (***Table 1***). Podocyte count error was bounded by one podocyte per glomerulus (median 0.61, ***Table 2***).

**Table 2:**
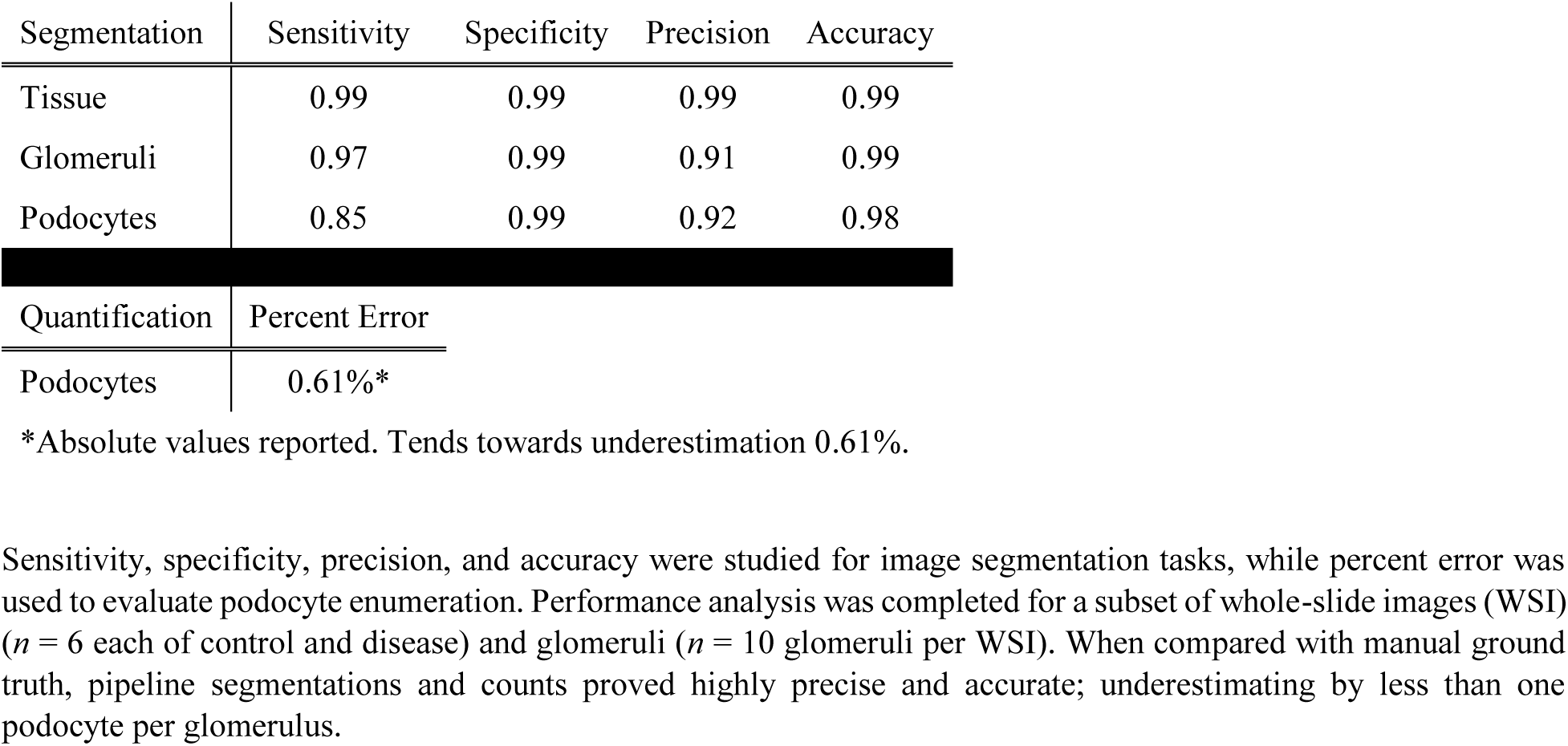
Computational performance analysis revealed highly precise and accurate podocyte enumeration as well as image segmentation.

| Segmentation | Sensitivity | Specificity | Precision | Accuracy |
| --- | --- | --- | --- | --- |
| Tissue | 0.99 | 0.99 | 0.99 | 0.99 |
| Glomeruli | 0.97 | 0.99 | 0.91 | 0.99 |
| Podocytes | 0.85 | 0.99 | 0.92 | 0.98 |
| Quantification | Percent Error |  |  |  |
| Podocytes | 0.61%* |  |  |  |

\*Absolute values reported. Tends towards underestimation 0.61%.
Sensitivity, specificity, precision, and accuracy were studied for image segmentation tasks, while percent error was used to evaluate podocyte enumeration. Performance analysis was completed for a subset of whole-slide images (WSI) ( $n = 6$ each of control and disease) and glomeruli ( $n = 10$ glomeruli per WSI). When compared with manual ground truth, pipeline segmentations and counts proved highly precise and accurate; underestimating by less than one podocyte per glomerulus.

### 3.3 Podocyte and glomerulus feature significance across murine models

Feature-based comparison of podocyte and glomerulus histomorphology was computed to determine whether quantified image features distinguished diseased tissue from normal tissue. Statistical analysis focused on the following image features: podocyte count, glomerulus area, glomerular podocyte density, total podocyte area, and glomerular podocyte coverage. As described in Methods, glomerular podocyte density and glomerular podocyte coverage are equivalent to the ratios of podocyte count: glomerular area and total podocyte area: glomerular area, respectively.

#### 3.3a Mouse-level disease indicators

Feature significance was first evaluated at the mouse level irrespective of biological sex (***Fig 5***). For each murine cohort, whole-slide podocyte and glomerulus features were compared between wild-type and disease using two sample *t*-tests (***Table 3***). Populations were defined by each mouse’s mean feature values. No feature proved significant for the T2DM A or HIVAN models. In contrast, histologic image features were significant indicators of disease across the T2DM B, Aging, FSGS (SAND), and Progeroid (*Ercc1^-/Δ^*) models. Glomerular podocyte density was the lead indicator in the Aging and FSGS (SAND) models, followed by glomerular area and glomerular podocyte coverage. Podocyte count was also significant in the FSGS (SAND) model. In the T2DM B model, podocyte count, glomerular area, glomerular podocyte density and total podocyte area were highly significant; glomerular podocyte coverage was not. Podocyte count and glomerular area were equally and highly indicative of the accelerated aging phenotype observed in *Ercc1^-/Δ^* Progeroid mice. Collectively, these results emphasize both podocyte and glomerulus morphometrics as strong indicators of disease.

**Figure 5:**
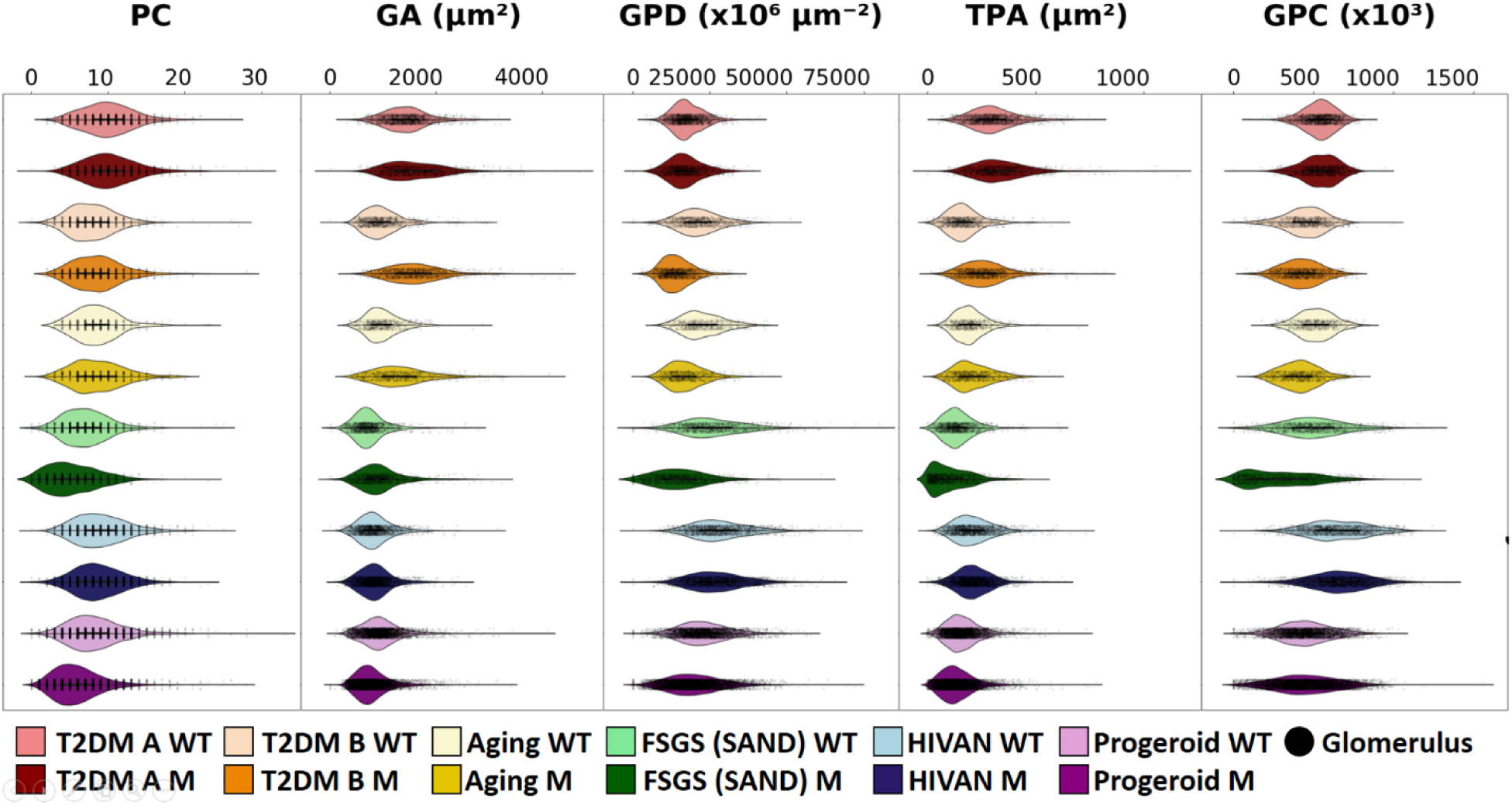
Podocyte and glomerular morphometrics for wild-type mice and disease phenotypes at the mouse level. *Legend.* T2DM A, Type-II diabetes mellitus A, the db/db model; T2DM B, Type-II diabetes mellitus B, the KKAy model; Aging, modeled after the NIA agent rodent colony; FSGS (SAND), the model of post-adaptive FSGS driven by SAND treatment; HIVAN, HIV-associated Nephropathy, the Tg26 model of collapsing glomerulopathy often classified as a form of FSGS; Progeroid, the accelerated aging model driven by *Ercc1^-/Δ^*. Violin plots showing measurements included glomerular podocyte count (PC), glomerulus area (GA) glomerular podocyte density (GPD), total podocyte area (TPA), and glomerular podocyte coverage (GPC). These measurements were compared across murine models. PC is defined as the number of podocyte nuclei per glomerular image. GA is the area of the glomerulus cross-section. TPA is computed as the cumulative area of labeled podocyte nuclei per glomerulus cross-section. GPD is computed as the ratio of absolute podocyte count to glomerulus cross-sectional area. Similarly, GPC is computed as the ratio of total podocyte area to glomerulus cross-sectional area. All metrics were converted to microns to facilitate biological interpretation. Plots illustrate the distribution of feature values across disease models with each black dot corresponding to a single glomerulus or data point.

**Table 3:** Podocyte and glomerulus morphometrics differentiated wild-type and disease phenotypes at the mouse level.

| *n* | *T2DM A* |  | *T2DM B* |  | *Aging* |  | *FSGS (SAND)* |  | *HIVAN* |  | *Progeroid* |  |
| --- | --- | --- | --- | --- | --- | --- | --- | --- | --- | --- | --- | --- |
|  | db/m | db/db | C57BL/6 | KKAy | Young | Old | No Tx | SAND | WT | Tg26 | WT | *Ercc1*<sup>-Δ</sup> |
|  | 6 | 4 | 10 | 10 | 6 | 5 | 11 | 8 | 11 | 13 | 20 | 31 |
| PC | 0.188 |  | 0.001** |  | 0.739 |  | 0.036* |  | 0.165 |  | <0.001*** |  |
| GA | 0.602 |  | <0.001*** |  | 0.005** |  | 0.002** |  | 0.842 |  | <0.001*** |  |
| GPD | 0.222 |  | <0.001*** |  | <0.001*** |  | <0.001*** |  | 0.495 |  | 0.061 |  |
| TPA | 0.911 |  | <0.001*** |  | 0.363 |  | 0.198 |  | 0.749 |  | 0.055 |  |
| GPC | 0.474 |  | 0.303 |  | 0.002** |  | 0.006** |  | 0.954 |  | 0.745 |  |

T2DM, type-2 diabetes mellitus; WT, wild-type; *n*, number of mice.
Glomerular podocyte count, PC; glomerulus area, GA; glomerular podocyte density, GPD; total podocyte area, TPA; glomerular podocyte coverage, GPC.
For each mouse model, pipeline-computed features were ranked based on their ability to differentiate between wild-type and diseased mice. In the T2DM B, Aging, FSGS (SAND), and Progeroid models, image features were significant indicators of disease. Podocyte morphometrics were of equal or greater significance than glomerular features. For those models wherein image features proved insignificant, statistical analysis would be reduplicated at the glomerulus level; no features were significant in the HIVAN model.
Statistical conclusions are based on two-sample *t*-tests comparing population means at a significance level of 0.05. Asterisks flag levels of significance: $p < 0.05^*$ , $p < 0.01^{**}$ , $p < 0.001^{***}$ .

#### 3.3b Glomerulus-level disease indicators

In Section 3.3a, feature values were computed at the mouse level by quantifying the mean feature value across all glomeruli derived from the respective mouse’s kidney section. Should disease manifest in a small subset of glomeruli, computation of the mean feature value may average out informative histopathologic phenomena. Therefore, comparison between wild-type and disease phenotypes was replicated at the glomerulus level, with test populations comprised of pooled wild-type and disease glomeruli across mice (***Table 4***). Similar feature significance was observed in the T2DM B and FSGS (SAND) models. In the Aging model, decreased glomerular podocyte counts were insignificant. While this result suggests an absence of podocyte depletion, old mice demonstrated a significant reduction in podocyte nuclear area, and thus this result might suggest a trend toward podocyte cell death, as nuclear condensation and shrinkage are fundamental apoptotic stages^8,51^. Glomerular area and glomerular area-driven features, glomerular podocyte density and glomerular podocyte coverage, were found to be significant. In the *Ercc1^-/Δ^* Progeroid mouse model, podocyte count, glomerular area, glomerular podocyte density, and total podocyte area were indicative of disease; glomerular podocyte coverage was not. Simultaneous reduction in glomerular area and total podocyte area resulted in an unremarkable difference in glomerular podocyte coverage between the Progeroid models’ *Ercc1^-/Δ^* and corresponding wild-type mice. Of particular interest were the T2DM A and HIVAN models. Feature values were found to be significant in these models at the glomerular level of analysis. Pooling of glomerular phenotypes across both the wild-type and mutant mice produced significantly different glomerulus populations. Significant reduction in glomerular podocyte count, glomerular podocyte density and glomerular podocyte coverage, as well as increased glomerular area, were observed in diabetes affected glomeruli. Unique to the HIVAN model was reduction in podocyte count and glomerular podocyte density; in the absence of increased glomerular area, podocyte loss drove glomerular podocyte density reduction.

**Table 4:** Podocyte and glomerulus morphometrics universally differentiate wild-type and disease populations at the glomerulus level.

| *n* | *T2DM A* |  | *T2DM B* |  | *Aging* |  | *FSGS (SAND)* |  | *HIVAN* |  | *Progeroid* |  |
| --- | --- | --- | --- | --- | --- | --- | --- | --- | --- | --- | --- | --- |
|  | db/m | db/db | C57BL/6 | KKAy | Young | Old | No Tx | SAND | WT | Tg26 | WT | *Ercc1*<sup>-Δ</sup> |
|  | 1396 | 958 | 773 | 970 | 527 | 603 | 1068 | 808 | 648 | 656 | 3788 | 5670 |
| PC | 0.007** |  | <0.001*** |  | 0.439 |  | <0.001*** |  | 0.002** |  | <0.001*** |  |
| GA | <0.001*** |  | <0.001*** |  | <0.001*** |  | <0.001*** |  | 0.650 |  | <0.001*** |  |
| GPD | <0.001*** |  | <0.001*** |  | <0.001*** |  | <0.001*** |  | <0.001*** |  | <0.001*** |  |
| TPA | 0.974 |  | <0.001*** |  | 0.007** |  | <0.001*** |  | 0.390 |  | <0.001*** |  |
| GPC | <0.001*** |  | 0.001** |  | <0.001*** |  | <0.001*** |  | 0.198 |  | 0.736 |  |

T2DM, type-2 diabetes mellitus; WT, wild-type; *n*, number of mice.
Glomerular podocyte count, PC; glomerulus area, GA; glomerular podocyte density, GPD; total podocyte area, TPA; glomerular podocyte coverage, GPC.
For those models wherein image features proved insignificant, statistical analysis was reduplicated at the glomerulus level. Pipeline-computed features were ranked based on their ability to differentiate between wild-type and diseased glomeruli in each model. Across all models, image features were significant indicators of disease. Podocyte morphometrics were of equal or greater significance than glomerular features. Notably, in the HIVAN model, podocyte metrics were significant while GA was not.
Statistical conclusions are based on two-sample *t*-tests comparing population means at a significance level of 0.05. Asterisks flag levels of significance: $p < 0.05^*$ , $p < 0.01^{**}$ , $p < 0.001^{***}$ .

#### 3.3c Mouse-level histologic indicators of proteinuria

Image features were correlated with terminal UACR to understand the relationship between renal micro anatomical integrity and functional outcome (proteinuria onset)^52–56^. Parametric Pearson correlation analysis was performed, comparing mice’s mean feature values with UACR measurements collected at the time of sacrifice. Correlation coefficients highlighted the predictive capacity of select features as well as each features’ relationship to proteinuria (increased UACR) (***Table 5***). Analysis was limited to those cohort studies wherein UACR was recorded: T2DM A, T2DM B, FSGS (SAND), and HIVAN. Significant relationships between histologic image features and terminal UACR were not observed in the T2DM A cohort. As mentioned before, the absence of an overt trend is attributed to the mild pathology characteristic of db/db mice^57–59^ in the T2DM A model. A subset of quantified image features was found to be predictive of proteinuria in the FSGS (SAND), HIVAN, and T2DM B models. The superior predictive capacity of glomerular podocyte density is attributed to the fact that this feature takes into account both absolute podocyte count and glomerular area. The observed shift in feature significance (*e.g.,* glomerular podocyte count in HIVAN) upon integration with clinical metrics warranted further exploration. Later analysis with respect to biological sex revealed a dichotomy in murine phenotype underlying feature patterns.

**Table 5:** Significant correlations were observed between histological image features and murine urinary albumin creatinine ratio (UACR) at time of sacrifice.

| *n* | *T2DM A* |  | *T2DM B* |  | *FSGS (SAND)* |  | *HIVAN* |  |
| --- | --- | --- | --- | --- | --- | --- | --- | --- |
|  | db/m | db/db | C57BL/6 | KKAy | No Tx | SAND | WT | Tg26 |
|  | 6 | 4 | 10 | 10 | 11 | 8 | 11 | 13 |
|  | *R*<sup>2</sup> | *p*-value | *R*<sup>2</sup> | *p*-value | *R*<sup>2</sup> | *p*-value | *R*<sup>2</sup> | *p*-value |
| PC | -0.481 | 0.159 | 0.670 | 0.002** | -0.465 | 0.052 | -0.344 | 0.108 |
| GA | 0.402 | 0.249 | 0.856 | <0.001*** | 0.643 | 0.004** | 0.533 | 0.009** |
| GPD | -0.537 | 0.110 | -0.809 | <0.001*** | -0.765 | <0.001*** | -0.539 | 0.008** |
| TPA | 0.173 | 0.632 | 0.827 | <0.001*** | -0.353 | 0.151 | 0.017 | 0.940 |
| GPC | -0.296 | 0.407 | -0.256 | 0.305 | -0.659 | 0.003** | -0.312 | 0.148 |

T2DM, type-2 diabetes mellitus; No Tx, no SAND treatment; WT, wild-type; *n*, number of mice.
Glomerular podocyte count, PC; glomerulus area, GA; glomerular podocyte density, GPD; total podocyte area, TPA; glomerular podocyte coverage, GPC.
UACR measurements were available for four of six mouse models: T2DM A, T2DM B, FSGS (SAND), and HIVAN. In the latter three models, select podocyte and glomerulus image features were highly correlated with UACR. More specifically, a reduction in GPD and GPC, as well as increase in GA, were significantly correlated with UACR increase in mice.
Correlation results between PC, GA, GPD, TPA, GPC and UAR were evaluated with parametric Pearson analysis (*R*<sup>2</sup> and *p* values reported). Values used for correlative analysis are pooled from assessment of renal microanatomical integrity at time of sacrifice and the corresponding functional outcome (UACR) at time of sacrifice. Asterisks flag levels of significance: *p* < 0.05\*, *p* < 0.01\*\*, *p* < 0.001\*\*\*.

### 3.4 Sex-associated feature significance across murine models

To assess whether disease manifestation was modulated by biological sex, feature-based comparison of murine histomorphology was refined to the sex level. For those cohorts including both male and female mice – FSGS (SAND), HIVAN, and Progeroid (*Ercc1^-/Δ^*) – whole-slide podocyte and glomerulus features were compared between wild-type and disease using two sample *t*-tests.

#### 3.4a Mouse-level disease indicators with respect to biological sex

We performed evaluation of feature significance at the mouse level (***Fig 6***). Due to sample size limitations and the requirements of statistical tests, this study was conducted for HIVAN and *Ercc1^-/Δ^* Progeroid mouse models only. For each of these murine cohort’s male and female populations, whole-slide podocyte and glomerulus features were compared between wild-type and disease using two sample *t*-tests (***Table 6***). Progeroid No feature was found to be significant in the HIVAN cohort. Of greater significance were observations from the *Ercc1^-/Δ^* Progeroid cohort wherein glomerular area was of equal significance in both sexes, while reduced podocyte count highly indicative of disease in males compared to females.

**Figure 6:**
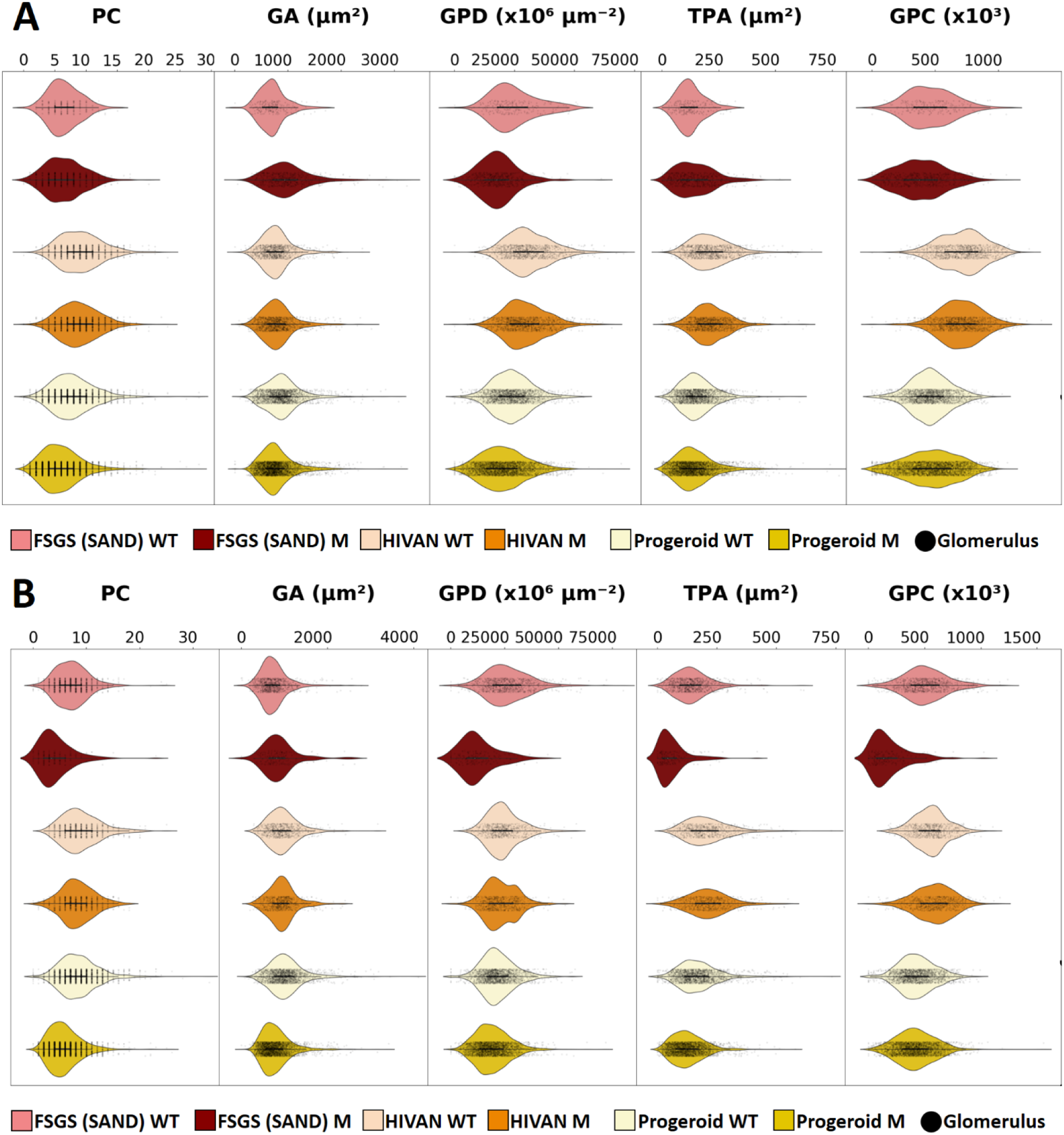
Female (A) and male mice (B) each demonstrate distinct podocyte morphometrics in a subset of models. Glomerular podocyte count, PC; glomerulus area, GA; glomerular podocyte density, GPD; total podocyte area, TPA; glomerular podocyte coverage, GPC. *Legend*. Data are shown as violin plots for wild-type controls (WT) and three glomerular disease models (M). The FSGS (SAND), HIVAN, and Progeroid (*Ercc1^-/Δ^*) models in this study included both male and female mice, and thus statistical analysis was completed to assess whether podocyte depletion was unique to female or male mice in each model. A) Plots illustrate the distribution of feature values across females in each disease model. B) Plots illustrate the distribution of feature values across males in each disease model. Each black dot corresponds to a single glomerulus or data point.

**Table 6:** Male and female mice demonstrate distinct podocyte morphometrics in disease.

| *n* | *Male* |  |  |  | *Female* |  |  |  |
| --- | --- | --- | --- | --- | --- | --- | --- | --- |
|  | *HIVAN* |  | *Progeroid* |  | *HIVAN* |  | *Progeroid* |  |
|  | WT | Tg26 | WT | *Ercc1*<sup>-Δ</sup> | WT | Tg26 | WT | *Ercc1*<sup>-Δ</sup> |
|  | 5 | 5 | 11 | 11 | 6 | 8 | 9 | 19 |
| PC | 0.177 |  | <0.001*** |  | 0.541 |  | 0.012* |  |
| GA | 0.531 |  | <0.001*** |  | 0.250 |  | 0.001** |  |
| GPD | 0.389 |  | 0.083 |  | 0.216 |  | 0.173 |  |
| TPA | 0.741 |  | 0.023* |  | 0.899 |  | 0.522 |  |
| GPC | 0.947 |  | 0.829 |  | 0.658 |  | 0.811 |  |

WT, wild-type; *n*, number of mice.
Glomerular podocyte count, PC; glomerulus area, GA; glomerular podocyte density, GPD; total podocyte area, TPA; glomerular podocyte coverage, GPC.
For those cohorts containing both biological sexes, feature significance was compared between male and female populations. A) HIVAN: No features were significant on the sex level. B) Progeroid: Reduction in PC was highly significant in male mice. Increase in GA was significant for both male and female mice.
Statistical conclusions are based on two-sample *t*-tests comparing population means at a significance level of 0.05. Asterisks flag levels of significance: $p < 0.05^*$ , $p < 0.01^{**}$ , $p < 0.001^{***}$ .

#### 3.4b Glomerulus-level disease indicators with respect to biological sex

To expand upon mouse-level observations, feature analysis for males and females was replicated at the glomerulus level (***Table 7***). In the FSGS (SAND) cohort, significant podocyte loss was unique to males. Further, podocyte loss was a much stronger indicator of disease in FSGS (SAND) males compared to the full cohort. Sex-based disparity in podocyte count also explains the diluted significance of podocyte count in the FSGS (SAND) correlational analysis. Similar dichotomy was observed in the HIVAN cohort. Male-derived glomeruli featured significant podocyte reduction, while female-derived glomeruli were larger on average. Although reduction in glomerular podocyte density was significant for both HIVAN-affected males and females, the underlying mechanism appears distinct. Reduction in glomerular podocyte density is driven by podocyte loss in males and glomerulomegaly in females. Such trends were not observed in the *Ercc1^-/Δ^* Progeroid cohort, wherein feature significance was uniform across mice, irrespective of sex.

**Table 7:** Glomeruli derived from male and female mice demonstrate distinct podocyte morphometrics in disease.

| *n* | *Male* |  |  |  |  |  | *Female* |  |  |  |  |  |
| --- | --- | --- | --- | --- | --- | --- | --- | --- | --- | --- | --- | --- |
|  | *FSGS (SAND)* |  | *HIVAN* |  | *Progeroid* |  | *FSGS (SAND)* |  | *HIVAN* |  | *Progeroid* |  |
|  | No Tx | SAND | WT | Tg26 | WT | *Ercc1*<sup>-Δ</sup> | No Tx | SAND | WT | Tg26 | WT | *Ercc1*<sup>-Δ</sup> |
|  | 850 | 172 | 593 | 638 | 1815 | 2597 | 218 | 496 | 932 | 1066 | 1973 | 3073 |
| PC | <0.001*** |  | 0.001** |  | <0.001*** |  | 0.548 |  | 0.162 |  | <0.001*** |  |
| GA | <0.001*** |  | 0.325 |  | <0.001*** |  | <0.001*** |  | 0.069 |  | <0.001*** |  |
| GPD | <0.001*** |  | 0.003** |  | <0.001*** |  | <0.001*** |  | <0.001*** |  | <0.001*** |  |
| TPA | <0.001*** |  | 0.271 |  | <0.001*** |  | <0.001*** |  | 0.851 |  | <0.001*** |  |
| GPC | <0.001*** |  | 0.001** |  | 0.030* |  | <0.001*** |  | 0.304 |  | 0.047* |  |

No Tx, no SAND treatment; WT, wild-type; *n*, number of glomeruli.
Glomerular podocyte count, PC; glomerulus area, GA; glomerular podocyte density, GPD; total podocyte area, TPA; glomerular podocyte coverage, GPC.
Feature significance was evaluated at the glomerulus level for male as well as female mice. Image features were ranked based on their ability to differentiate between wild-type and diseased glomeruli in each model. Across all models, image features were significant indicators of disease. Podocyte morphometrics were of equal or greater significance than glomerular features. A) FSGS (SAND) and HIVAN: PC was a significant indicator of disease in males exclusively. B) Progeroid: All features were significant in both male and female derived glomeruli.
Statistical conclusions are based on two-sample *t*-tests comparing population means at a significance level of 0.05. Asterisks flag levels of significance: $p < 0.05^*$ , $p < 0.01^{**}$ , $p < 0.001^{***}$ .

#### 3.4c Mouse-level histologic indicators of proteinuria with respect to biological sex

To assess whether the observed histologic dichotomies were mirrored in functional outcomes, our prior correlational analysis was refined to the sex-level. Only those cohort studies including both male and females as well as terminal UACR measurements were studied. These included the FSGS (SAND) and HIVAN models only. Parametric Pearson correlation analysis was computed as before with reportage of correlation coefficients and corresponding *p*-values (***Table 8***). In the HIVAN cohort, no correlation was observed between image features and terminal UACR. As for the FSGS (SAND) cohort, the dichotomy observed between male and female glomerulus populations was mirrored in the current correlational analysis. Reduction in podocyte nuclear count was correlated with proteinuria in male FSGS (SAND) mice exclusively. Furthermore, while glomerular podocyte density was correlated with proteinuria in both males and females, the strength and significance of correlation was much greater in males. All podocyte-related features were predictive of proteinuria in male mice, while glomerular area was not. This observation emphasizes the role of podocyte loss in reduced glomerular podocyte density and glomerular podocyte coverage, and potentiates these features as image-based estimates for glomerular filtration barrier integrity and proteinuria onset^7–9,60–62^.

**Table 8:** Significant correlations were observed between histological image features and murine urinary albumin creatinine ratio (UACR) at time of sacrifice.

| *n* | *Male* |  |  |  | *Female* |  |  |  |
| --- | --- | --- | --- | --- | --- | --- | --- | --- |
|  | *FSGS (SAND)* |  | *HIVAN* |  | *FSGS (SAND)* |  | *HIVAN* |  |
|  | No Tx | SAND | WT | Tg26 | No Tx | SAND | WT | Tg26 |
|  | 9 | 2 | 5 | 5 | 2 | 5 | 6 | 8 |
| | $R^2$ | *p*-value | $R^2$ | *p*-value | $R^2$ | *p*-value | $R^2$ | *p*-value |
| PC | -0.736 | 0.010* | -0.336 | 0.377 | -0.048 | 0.918 | -0.070 | 0.811 |
| GA | 0.463 | 0.152 | 0.488 | 0.182 | 0.695 | 0.083 | 0.476 | 0.085 |
| GPD | -0.780 | 0.005** | -0.544 | 0.130 | -0.763 | 0.046* | -0.373 | 0.189 |
| TPA | -0.684 | 0.020* | 0.108 | 0.781 | -0.055 | 0.905 | 0.039 | 0.894 |
| GPC | -0.747 | 0.008** | -0.101 | 0.796 | -0.552 | 0.199 | -0.147 | 0.617 |

No Tx, no SAND treatment; WT, wild-type; *n*, number of mice.
Glomerular podocyte count, PC; glomerulus area, GA; glomerular podocyte density, GPD; total podocyte area, TPA; glomerular podocyte coverage, GPC.
Sex-specific UACR measurements were available for two of six mouse models: FSGS (SAND) and HIVAN. A) FSGS (SAND): In male mice, podocyte morphometrics were highly correlated with UACR; GA was not. In female mice, GPD was correlated with UACR. B) HIVAN: No features were significantly correlated with UACR for either sex.
Correlation results between PC, GA, GPD, TPA, GPC and UACR were evaluated with parametric Pearson correlation analysis ( $R^2$ and *p* values reported). Values used for correlative analysis are pooled from assessment of renal microanatomical integrity at time of sacrifice and the corresponding functional outcome (UACR) at time of sacrifice. Asterisks flag levels of significance: $p < 0.05^*$ , $p < 0.01^{**}$ , $p < 0.001^{***}$ .

### 3.5 Patient-level feature significance in human DN

Biopsy level features were compared among DN subjects based on their Tervaert classification^33^ (***Fig S1***) as well as their outcome. Analysis of variance was used to evaluate feature significance across Tervaert stages (***Table 9***). *Post hoc* tests were then applied to identify the particular DN stage(s) (Table 10). Podocyte count was the lead indicator of DN stage at the patient-level, with overt podocyte loss defining the transition from DN stage IIb to III. Glomerular area was also significant, but to a lesser extent than podocyte-inclusive features (glomerular podocyte count and podocyte coverage, total podocyte area). Similar to podocyte count, reduction in total podocyte area and glomerular podocyte coverage characterized the transition from stage IIb to III. Meanwhile, pathologic increase in glomerular area differentiated stage IIb from IV. These observations are consistent with established histopathologic classification criteria^33,63^, discussed later.

**Table 9:** Podocyte and glomerulus morphometrics differentiated diabetic nephropathy stages at the patient level.

| Feature | $p$-value ($n = 45$) |
| --- | --- |
| PC | 0.005** |
| GA | 0.041* |
| GPD | 0.058 |
| TPA | 0.012* |
| GPC | 0.014* |

$n$ , number of patients.
Glomerular podocyte count, PC; glomerulus area, GA; glomerular podocyte density, GPD; total podocyte area, TPA; glomerular podocyte coverage, GPC.
Pipeline-computed features were ranked based on their ability to differentiate between diabetic nephropathy stages defined by the Tervaert classification scheme. PC was the most significant indicator of disease in diabetic nephropathy, followed by TPA and GPC; GA was the least significant indicator.
Statistical conclusions are based on one-way ANOVA comparing population means at a significance level of 0.05. $H_0$ : At least one population mean is different. Asterisks flag levels of significance: $p < 0.05^*$ , $p < 0.01^{**}$ , $p < 0.001^{***}$ .

**Table 10:** Significant differences in podocyte and glomerulus morphometrics are consistently observed between diabetic nephropathy (DN) stages IIb and III.

| Feature | Significantly different diabetic nephropathy stages | $p$-value ($n = 45$) |
| --- | --- | --- |
| PC | IIb from III | 0.005** |
| GA | IIb from IV | 0.041* |
| GPD | III from IIa, IIb, and IV | 0.058 |
| TPA | IIb from III | 0.012* |
| GPC | IIb from III | 0.014* |

$n$ , number of patients.
Glomerular podocyte count, PC; glomerulus area, GA; glomerular podocyte density, GPD; total podocyte area, TPA; glomerular podocyte coverage, GPC.
Pairwise tests were completed to identify which DN stages underlie significant differences in patient feature data. Asterisks flag levels of significance: $p < 0.05^*$ , $p < 0.01^{**}$ , $p < 0.001^{***}$ .

### 3.6 Prediction of ESKD in DN

To assess the predictive power of histological image features, a series of binary logistic regression models were evaluated for patient progression to ESKD. Image features functioned as explanatory variables, while ESKD occurrence defined the response (***Fig 7***). A model was fit for each feature-outcome combination (e.g., glomerular podocyte count as the variable and ESKD as the response). All models were assessed using (i) the Wald Chi-Squared Test to assess the statistical significance of the explanatory variable, and (ii) the Deviance Goodness-of-Fit Test to determine whether or not the model fit the data well. Measures of model performance as well as *p*-values were recorded for subsequent feature ranking (***Table 11***). Once again, statistical analysis focused on the following image features: podocyte count, glomerular area, glomerular podocyte density, total podocyte area, and glomerular podocyte coverage. A model was also fit using patients’ eGFRs at time of biopsy, as eGFR is known to be a predictor of ESKD in DN^64,65^. Quantified image features, excluding glomerular podocyte density, and eGFR were found to be predictive of outcome with *p*-value < 0.05. This result underscores the potential for computational histological image analysis in the assessment and prognostication of DN.

**Figure 7:**
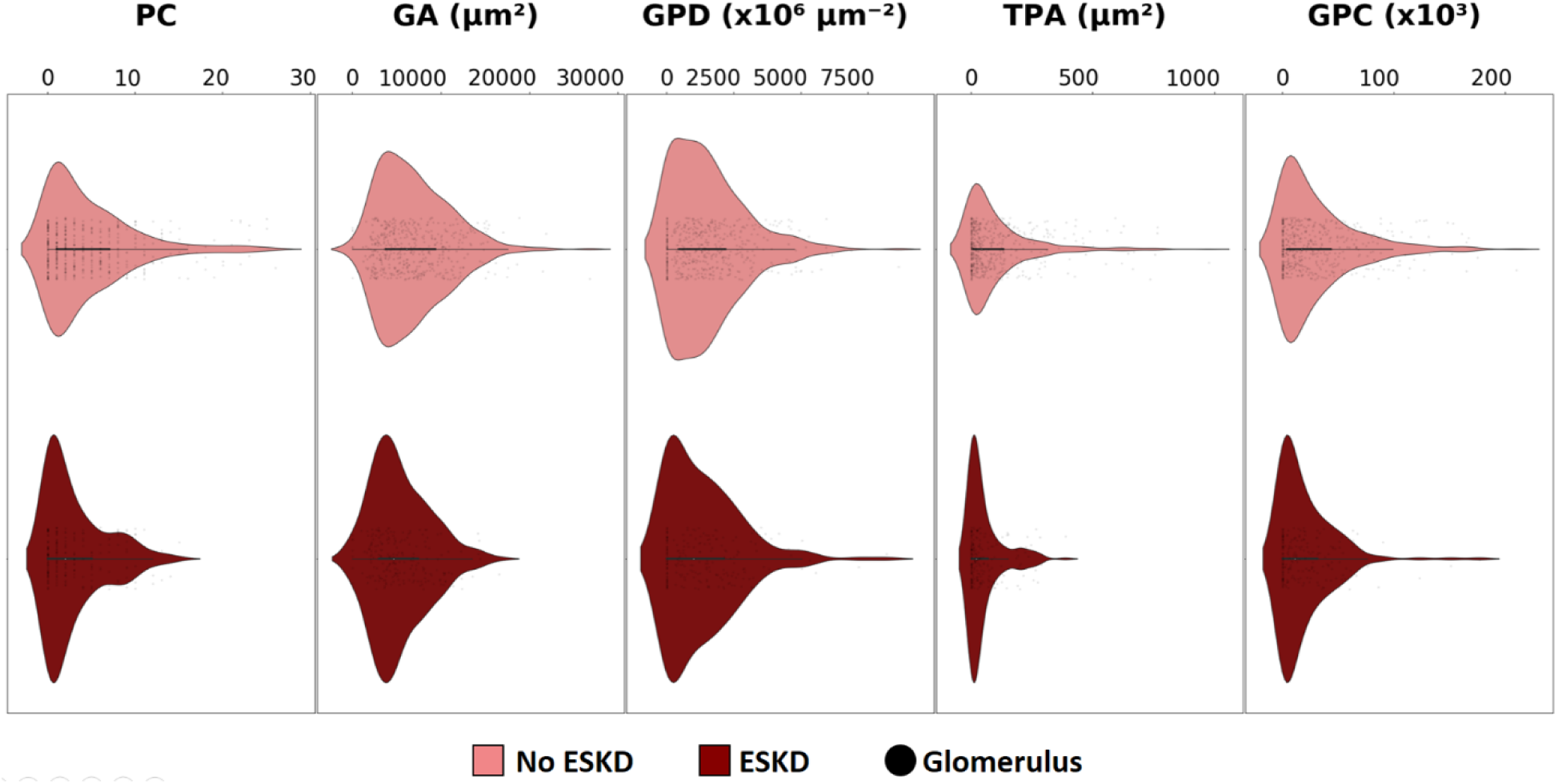
Glomeruli derived from Diabetic Nephropathy (DN) kidney biopsies demonstrate distinct podocyte and glomerulus morphometrics based on patient outcome. Glomerular podocyte count, PC; glomerulus area, GA; glomerular podocyte density, GPD; total podocyte area, TPA; glomerular podocyte coverage, GPC. The DN cohort featured patients that did and did not progress to ESKD. Plots illustrate the distribution of feature values for both outcomes with each black dot corresponding to a single glomerulus or data point.

**Table 11:** Podocyte morphometrics are significant predictors of patient outcome in Diabetic Nephropathy (DN).

| Variable | Coefficient | 95% CI | *p*-value of Wald test | Chi-square score of model fit (*df* = 1) | *p*-value of Chi-square score |
| --- | --- | --- | --- | --- | --- |
| PC | -0.503 | [0.337, 0.970] | 0.037* | 7.749 | 0.005** |
| GA | -0.001 | [0.999, 0.999] | 0.024* | 7.631 | 0.006** |
| GPD | -0.001 | [0.998, 1.001] | 0.1628 | 2.139 | 0.144 |
| TPA | -0.024 | [0.957, 0.997] | 0.023* | 10.325 | 0.001** |
| GPC | -0.063 | [0.887, 0.994] | 0.031* | 6.969 | 0.008** |
| eGFR | 0.303 | [1.017, 1.802] | 0.038* | 5.011 | 0.025* |

eGFR, estimated glomerular filtration rate at time of biopsy.
Glomerular podocyte count, PC; glomerulus area, GA; glomerular podocyte density, GPD; total podocyte area, TPA; glomerular podocyte coverage, GPC.
Binary logistic regression models were evaluated for feature-based prediction of patient outcome in diabetic nephropathy. Each pipeline-computed image feature was evaluated as the explanatory variable in prediction of ESKD incidence (response variable). Response frequency was fourteen out of forty-five patients. Coefficients, *p*-values and confidence intervals reported. Asterisks flag levels of significance: $p < 0.05^*$ , $p < 0.01^{**}$ , $p < 0.001^{***}$ .

### 3.7 Disbursement of *PodoCount* via the cloud

PodoCount was deployed as a cloud-based plugin on the Sarder Lab’s Digital Slide Archive (DSA)^66^ (***Fig 8***). This seamless integration was facilitated by *HistomicsTK*, a web-based tool that (i) allows for installation of user-defined algorithms as plugins in a virtual user-interface (UI), *HistomicsUI*, and (ii) is supported by the *OpenSlide*^67^ library for handling proprietary digital pathology WSI formats. These features render our *PodoCount* plugin as a universally accessible podocyte quantification application for all users, irrespective of their operating system or coding experience. To quantify podocytes in their image data, users (i) upload a WSI of IHC-labeled kidney specimen and the corresponding glomerulus annotation file to the DSA, (ii) select the option for the *PodoCount* tool, and (iii) download the output image feature and podocyte annotation files. Features are summarized in .csv (comma-separated values) format as Microsoft Excel files, while podocyte annotations are prepared as xml files compatible with standard desktop pathology viewers (*e.g*., Aperio ImageScope). To preview output annotations in the cloud, users need to only select the option “TranslateXMLToJson” included as a part of the plugin. This function automatically converts the annotation file into a web-format, displaying green podocyte nuclear contours in the web-viewer (***Fig 8***). To generate a glomerulus annotation file for input to *PodoCount*, users may apply the glomerulus detection plugin developed by *Lutnick et al*.^68,69^, readily available via the DSA.

**Figure 8:**
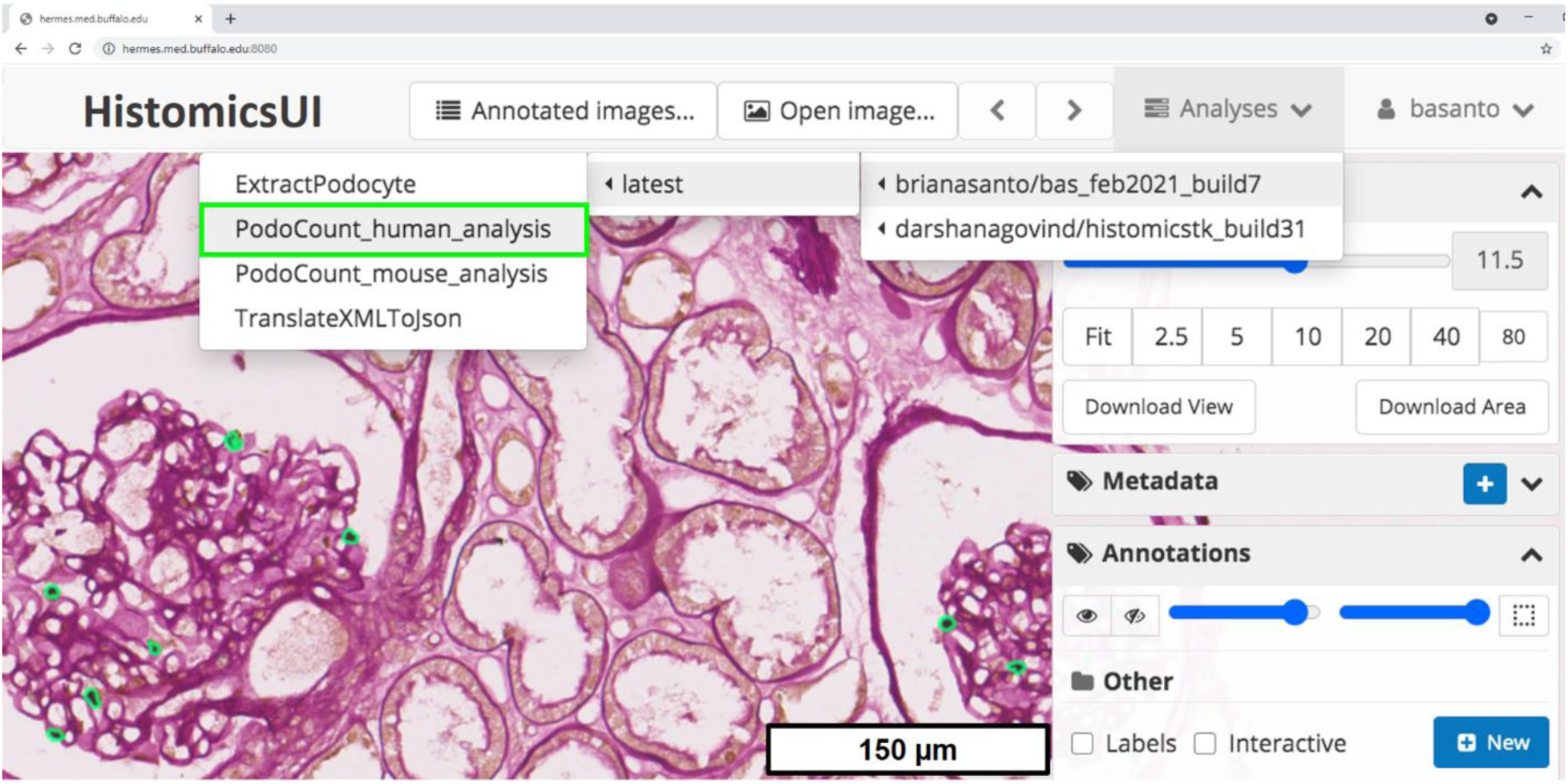
Our tool was converted into a cloud-based plug-in to provide for user-friendly, universally accessible podocyte quantification. The described podocyte quantification pipeline was converted into a web plug-in for our lab’s cloud-based WSI archiver and viewer. Upon upload of digitized kidney histopathology, users may apply our *PodoCount* (bright green) plug-in for whole-slide podocyte nuclear enumeration and morphometric analysis. Quantified morphometrics are output as *Microsoft Excel* feature files for user download. Whole-slide podocyte annotation files compatible with standard desktop digital pathology viewers (e.g., Aperio ImageScope) are also output. Figure features a diabetic nephropathy stage IIb kidney biopsy.

## 4. DISCUSSION

In this work we introduce *PodoCount*, a novel tool for automated, whole-slide assessment of podocyte depletion. This computational tool is the first of its kind, enabling quantification of podocyte depletion from a holistic, big-data perspective. These claims are supported by the comprehensive cohort study described herein. We validated this new method in a complex dataset comprising male and female-derived, digital kidney specimens from six distinct mouse models (*n* = 135 mice) as well as human diabetic nephropathy (*n* = 45 patients). These data were curated from multiple institutions and feature the highly variable stain, image, and tissue quality hindering development of generalizable computational frameworks^19–21,23,70^. *PodoCount* navigated this challenging dataset well, achieving highly precise and accurate segmentation as well as enumeration of podocyte nuclei. When paired with strategic feature engineering, this computational performance facilitated robust quantification of absolute and relative podocyte depletion across renal pathologies. Before delving into the unique podocyte representations, or physical image attributes, observed in this study, key details of our methodology must be addressed within the context of extant podocyte analytics. More specifically, our combination of biological and engineering technique informed the choice of (i) podocyte marker, and (ii) nuclear-based quantification from two-dimensional (2D) cross sections. We assert and will discuss the validity of these decisions toward scientific reproducibility and feasibility.

### 4.1 p57^kip2^ is a robust podocyte label

Four cell types contribute to the structure and function of the renal glomerulus: podocytes, parietal epithelial cells (PECs), glomerular capillary endothelial cells, and mesangial cells. Residing in close proximity, these cell types are immensely difficult to resolve in traditional stains^10^. Quantification of each cell population requires labeling with antibodies directed at cell-specific targets. An optimal marker must exploit the unique characteristics of podocytes to achieve sensitive and specific detection. Hallmarks of podocyte phenotype include cellular quiescence and limited proliferative capacity^71,72^. This characteristic dormancy is promoted by *de novo* expression of p57^kip2^ – a cyclin dependent kinase inhibitor modulating cell cycle arrest and terminal differentiation^35^. Podocyte-exclusive expression of p57^kip2^ within the glomerular microenvironment render p57^kip2^ a better maker than protein targets actively expressed by both podocytes and PECs^9–11,73–75^. The evolving role of podocyte dedifferentiation and phenotypic transformation in glomerulopathy further complicates marker selection^8,76,77^. Podocyte-driven glomerular disease assessment requires analysis of functional integrity, and thus necessitates confirmation of the terminally-differentiated podocyte phenotype. To our knowledge, p57^kip2^ is the only marker widely validated in the literature for podocyte terminal differentiation^35^.

### 4.2 Podocyte nuclear representations are strong indicators of cellular pathology

Podocytes exemplify the structure-function relationship that transcends biological fields. Fine processes, representing cytoplasmic projections, delicately wrap about the glomerular capillaries to form a highly selective barrier at the filtration interface, influenced by charge and size. In disease states, these delicate processes fuse, trading off filtration integrity and fluid balance for cell survival, at the cost of allowing proteinuria and promoting glomerulosclerosis^8,60^. Based on this pathobiology, measuring cytoplasmic surface area seems a logical approach to assessing podocyte depletion. However, quantification of podocyte cytoplasmic area does not equate to glomerular podocyte number. The complex morphology of podocyte cytoplasm undermines the ability to individually assess highly interdigitated cells. This morphological intricacy also requires well-refined staining to ensure objective quantification of cytoplasm.

Popular podometric approaches have used immunofluorescence and state-of-the-art microscopy to identify and measure podocyte cytoplasm^12,13,78,79^. While innovative, some of these studies still rely on a nuclear label to (i) irrefutably identify a cell as a podocyte, and (ii) estimate podocyte depletion. Furthermore, several results suggest that nuclear-based podocyte quantification is more than sufficient. For example, in a transgenic mouse and a dose-dependent model of podocyte depletion, glomerular podocyte density – a nuclear-based feature – proved to be a lead indicator of podocyte depletion^13^. In addition, nuclear volume was confirmed as a positive indicator of podocyte hypertrophy^13^. Average podocyte nuclear volume increased dramatically in mild and moderately podocyte-depleted mice, compared to control. All the while, podocyte nuclear-to-cytoplasmic ratio was consistent across murine phenotypes. Podocytes scaled evenly in size, with increasingly significant correlation between nuclear and cytoplasmic volume observed for mice with progressive podocyte depletion^13^. Finally, podocytes are rarely binucleate, suggesting that enumeration of nuclear profiles provides an estimation of absolute count^80^. These observations suggest that podocyte nuclei provide a strong foundation for both computations of glomerular podocyte density as well as assessment of podocyte depletion.

### 4.3 2D nuclear-based quantification outperforms its 3D, whole-cell counterparts

Support for nuclear-based quantification of podocyte depletion is augmented by the need for feasibility. Dimensionality is an important consideration. While informative, 3D podocyte visualization is not essential for objective quantification of depletion. According to *Puelles et al*., podocyte enumeration in adjacent tissue sections can provide a discrepant number of podocytes per glomerulus^13^. However, this observation was based on a single glomerulus.

*PodoCount*, the computational pipeline that we describe here, analyzes whole murine kidney sections, computing podocyte depletion statistics from large glomerulus populations (*n* = 100 per kidney cross-section, on average)^4^. When comparing disease states, across a cohort of fifty (*e.g.,* the *Ercc1^-/Δ^* Progeroid mouse cohort), sample size quickly grows to *n* = 2500 glomeruli per murine phenotype. From probability theory and the law of large numbers, we know that as our glomerular sample population increases, quantified glomerular features will converge to the true, mean value^81,82^. Therefore, the ability of *PodoCount* to efficiently quantify podocyte depletion at large-scale favors optimal estimation of glomerular podocyte depletion, compared to less scalable 3D or other 2D methods considering serial tissue sections.

Prior studies report another key finding: the correlation of podometric estimates between 2D and 3D methods. Podocyte counts derived via 2D and 3D approaches are significantly correlated, as were nuclear-based estimates of glomerular podocyte density^13^. Irrespective of disease status, glomerular podocyte density was strongly and highly correlated between 2D and 3D methods (*R* = 0.94; *p* < 0.001)^13^. The same held true for cytoplasm-based quantification of depletion (*R* = 0.87; *p* < 0.001)^13^. Significant correlation, across features and disease phenotypes, raises a question as to whether 3D assessment is absolutely essential for podometric assessment of glomerular disease. High resolution 3D methods are costly, both with respect to time and resources. Optical sectioning requires access to state-of-the-art confocal microscopy systems as well as an abundance of time and money for fluorescence staining. Our method provides for efficient, large-scale podocyte analysis from IHC-labeled tissues, without introducing (i) the bias characteristic of common methods, (ii) the time-consuming, impracticality of gold-standard stereological techniques, or (iii) the costs of fluorescence-based methods. Successful incorporation of podometric analysis in experimental and clinical workflows is contingent upon a method that maximizes feasibility and reproducibility while also minimizing cost.

All of this being said, *PodoCount*’s ability to generalize across heterogeneous digital pathology datasets allows us to complete nuclear-based quantification of podocyte depletion from an unprecedented, big data perspective. *PodoCount* can be used to explore diverse biological hypotheses is not limited to podocytes. The tool sets the stage for ubiquitous quantification of intraglomerular cell populations from immuno-labeled nuclei. The infrastructure is already in place to study, for example, fluctuations in resident cell populations amidst glomerulomegaly^79^. Following such studies, our method could be expanded to quantify podocyte cytoplasm and thus whole podocytes.

### 4.4 Podocyte depletion as a ubiquitous indicator of renal disease

According to the podocyte depletion hypothesis, podocyte depletion may manifest (i) absolutely, as a reduction in glomerular podocyte count, or (ii) relatively, when pathologic increase in glomerular area reduces podocyte spatial density^3^. Both modes of depletion may coincide^78^, as observed in several of our cohorts (***Tables 3-4, 6-7***). Through our comprehensive study, we learned that podocyte metrics are reproducibly indicative of disease; even in the absence of benchmark histopathology (e.g., glomerulomegaly^63^). We also found that podocyte representations are unique to a disease model, and in some cases to biological sex.

Histologic image features evaluated for statistical significance in this work were glomerular podocyte (nuclear) count, glomerular area, glomerular podocyte density, total podocyte area, and glomerular podocyte coverage. On the mouse level, these features differentiated murine phenotypes across models. Namely, in our second diabetic model, T2DM B, glomerular podocyte count, glomerular area, glomerular podocyte density, and total podocyte area were equally significant indicators of disease. Both absolute and relative depletion are consistent with diabetic pathology^83^, as is increased glomerular area. Urinary albumin measured at time of sacrifice supports these results, with all four features strongly predictive of proteinuria. Similarly, in the Aging model, we observed reduced glomerular podocyte density, increased glomerular area, and thus relative podocyte depletion were characteristic of older mice. These findings are consistent with the pathology of aging nephrons^16,78^. Further, in the FSGS (SAND) model, both absolute and relative podocyte depletion were observed, with significant reduction in glomerular podocyte density, coverage, and count, as well as increased glomerular area. Glomerular podocyte coverage was markedly more significant than podocyte count. We believe that this disparity in significance, as well as the combination of absolute and relative depletion, is a product of biological sex. When we refined our analysis to the sex level and pooled glomeruli across mice, we found that absolute reduction in podocyte count was unique to male mice. UACR measured at time of sacrifice supports this conclusion, with podocyte count strongly predictive of proteinuria in males exclusively. The FSGS (SAND) model is designed to emulate post-adaptive FSGS. Given that the incidence and severity of FSGS is often greater in males, the FSGS (SAND) model and thus our podocyte metrics may reflect a pro-male phenotype. Moreover, in the *Ercc1^-/Δ^* Progeroid model, absolute podocyte depletion and increased glomerular area were identified as indicators of accelerated aging.

Significant features were not observed in the T2DM A or HIVAN models. Absence of feature significance was attributed to mild disease pathology in db/db or Tg26 mutants showing variable expressivity. We also believe that incomplete penetrance in a subset of mice undermined feature significance at the mouse level. When pooled across all mice of a given murine phenotype, wild-type and mutant glomerulus populations were significantly different for both models. Refinement of analysis to the sex level left findings unchanged. Unlike the FSGS (SAND) model, unique pathologies were not observed on the basis of sex. Expansion of our statistical analysis to the glomerulus level pointed to absolute podocyte depletion, with significant reduction in glomerular podocyte count and coverage, but no change in glomerular area. UACR measurements collected at time of sacrifice support the above conclusions, with a subset of HIV-afflicted mature mice failing to manifest proteinuria. Similar to the diabetic nephropathy study, discussed next, clinical measures of kidney function (e.g., UACR, eGFR) provided a ground truth or baseline for validation of murine phenotypes.

From the human DN cohort, we learned that podocyte image features are a valuable prognostic tool. Podocyte metrics successfully differentiated patients (*n* = 45) Tervaert classification, highlighting the transition from stage IIb to III as a key turning point in diabetic nephropathy pathology (***Table 10***). Intriguingly, *PodoCount* metrics predicted patient outcomes just as well as eGFR measured at time of biopsy – a clinical indicator of CKD^64^. Our engineered feature for total podocyte area proved to be the best predictor of patient progression to ESKD, when compared to eGFR and glomerular area. These findings underscore the prognostic power of podometrics and potentiate our *PodoCount* tool as a valuable addition to the clinical toolbox.

### 4.5 *PodoCount* transcends the limitations of current techniques

To ensure both universal accessibility and reproducibility, we established *PodoCount* in the cloud. As a cloud-based plugin to our lab’s DSA, any researcher or clinician can now study podocyte metrics independent of their coding experience or computer’s operating system. We have provided a direct, public link to our web-based plugin, as well as all of our data and codes (See *Data Availability*). Researchers, clinicians, and the simply curious, alike, may choose to experiment with our data or analyze their own. Further the shared docker image will allow anyone to reproducibly establish our developed tool in their own server as a web-plugin via *HistomicsUI* for end-users. *PodoCount* is to our knowledge the first podocyte analytic optimized across an internationally sourced, multi-institutional dataset, offering whole-slide podocyte quantification at the touch of a button. *PodoCount* transcends computational pathology barriers, exemplifying how biological and engineering expertise can produce a tool with unprecedented accessibility and generalizability that potentiates cloud-based analysis as an avenue for podometric standardization.

## AUTHOR CONTRIBUTIONS

BAS conceptualized and performed the quantitative analyses, designed and conducted the computational methods, interpreted the results, and wrote the manuscript. DG and DM assisted with the adaptation of *PodoCount* codes to the cloud format. PD assisted with completion of ground truth podocyte counts. XY assisted in manuscript preparation. T2DM A, B and Aging cohorts were generated by the Levi Lab (XXW, KM, BAJ, ML). The FSGS (SAND) and HIV mouse cohorts were generated by the Kopp Lab (JBK). JBK also assisted in manuscript editing. The *Ercc1^-/Δ^* Progeroid cohort and littermate controls were generated by the Niedernhofer Lab (LJN). HSS and MKC provided human biopsy and clinical data for diabetic patients. AR conceived and optimized the staining technique that enables this computational method. AR also assisted in manuscript preparation. PS conceived the idea of automated computational quantification and enumeration of podocytes from digital whole slide images in the domain of renal pathology, and making this tool available for nephrology community via an open-source cloud resource. PS also assisted in manuscript preparation, coordinated with the study team, assisted in study design, supervised the computational implementation, and critically analyzed the results.

## ACKNOWLEDGEMENTS

We are grateful to Brendon Lutnick, the developer of H-AI-L tool and the implementer of the DSA tool in the Sarder Lab, for his guidance. We acknowledge the assistance of the Reference Histology Core at Johns Hopkins University School of Medicine, Baltimore, Maryland. We acknowledge the assistance of the Seoul National University Hospital Human Biobank, a member of the National Biobank of Korea, which is supported by the Ministry of Health and Welfare, Republic of Korea, for provision of human bio specimens used.

