## Supplemental Figure and Table for "PodoCount: A robust, fully automated whole-slide podocyte quantification tool"

**Figure S1: Diabetic Nephropathy (DN) kidney biopsies feature distinct podocyte and glomerulus morphometrics across Tervaert classes.**

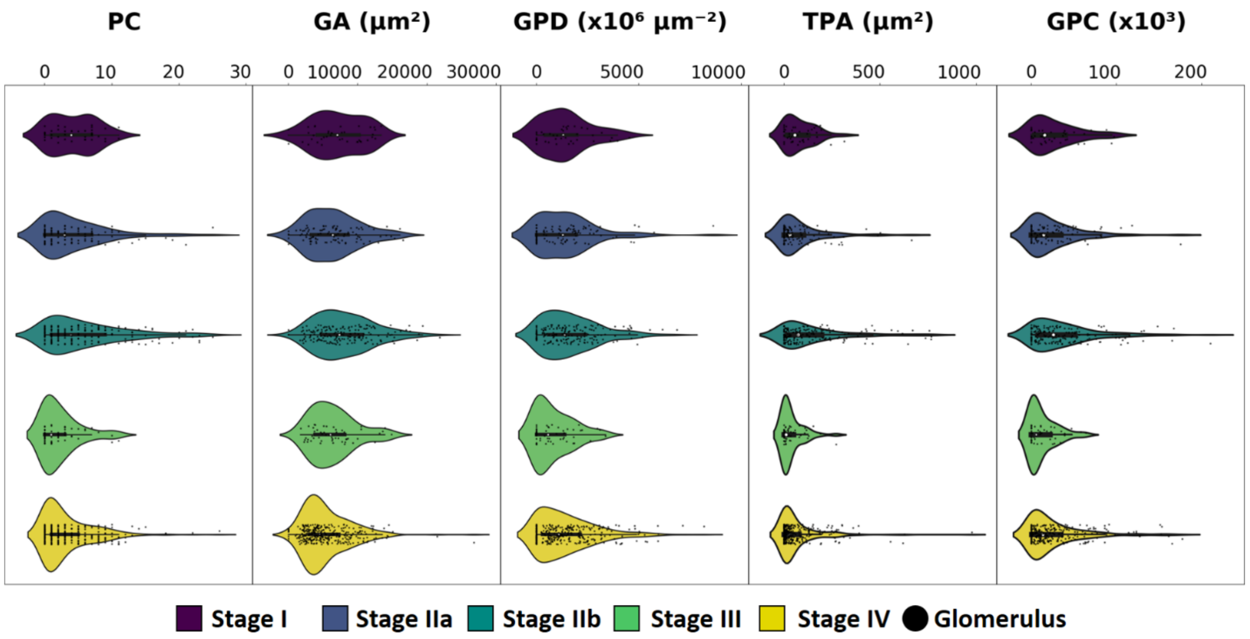


The diabetic nephropathy cohort included patients classified as stages I, IIa, IIb, III, and IV. Feature significance was compared between diabetic nephropathy stages. Plots illustrate the distribution of feature values across disease stages with each black dot corresponding to a single glomerulus or data point.

**Table S1: A comprehensive feature list including definitions is provided for ease of biological interpretation of computational findings.**

| Feature | Definition |
|---|---|
| PC | Also referred to as absolute podocyte count. The quantity of podocyte nuclei per glomerulus unit. Given that bi-nucleate podocytes are rare, nuclear enumeration provides an accurate estimation of glomerular podocyte count. |
| GA | The cross-sectional area of the glomerulus unit. |
| GPD | Also referred to as glomerular podocyte density. Computed as the ratio of podocyte count to glomerulus area. Approximates the spatial density of podocytes. |
| TPA | Also referred to as cumulative podocyte area. Computed as the cumulative area of podocyte nuclei for a given glomerulus. |
| GPC | Also referred to as glomerular podocyte coverage. Computed as the ratio of total podocyte nuclear area to glomerulus cross-sectional area. |
| PN Area | The cross-sectional area of a podocyte nucleus (PN). Computed as the total number of pixels comprising a segmented PN. |
| PN Bounding Box Area | The total number of pixels comprising the bounding box of a given PN. Here, the bounding box is the minimum or smallest enclosing box for a point set (PN pixels). |
| PN Convex Area | The total number of pixels comprising the convex hull of a given PN. Here, the convex hull is the smallest convex polygon enclosing the PN. |
| PN Eccentricity | The eccentricity of an ellipse fit to the PN. Here, the eccentricity is the ratio of the focal distance over the major axis length. Eccentricity ranges between zero and one, where a value of zero indicates a perfect circle. |
| PN Equivalent Diameter | The diameter of a circle with the same area as the PN. |
| PN Extent | The ratio of the PN's area to the PN's bounding box area. |
| PN Major Axis | The length of the major axis of the ellipse fit to the PN. |
| PN Minor Axis | The length of the minor axis of the ellipse fit to the PN. |
| PN Max Pixel Value | The maximum pixel value enclosed within the PN region. |
| PN Mean Pixel Value | The mean pixel value enclosed within the PN region. |
| PN Min Pixel Value | The minimum pixel value enclosed within the PN region. |
| PN Orientation | Given an ellipse fit to the PN, the angle between the ellipse's major axis and a horizontal line through the ellipse. |
| PN Perimeter | The number of pixels comprising the boundary of the PN. |
| PN Solidity | The ratio of the PN's area to the PN's convex area. |

PC, podocyte count; GA, glomerulus area, GPD, glomerular podocyte density; TPA, total podocyte area; GPC, glomerular podocyte coverage; PN, '…', podocyte nuclear '…'.

All podocyte nuclear features are recorded as the average value for each glomerular podocyte population.
